# Adaptive post-transcriptional reprogramming of metabolism limits response to targeted therapy in BRAF^V600^ melanoma

**DOI:** 10.1101/626952

**Authors:** Lorey K Smith, Tiffany Parmenter, Margarete Kleinschmidt, Eric P Kusnadi, Jian Kang, Claire A Martin, Peter Lau, Julie Lorent, Anna Trigos, Teresa Ward, Aparna D Rao, Emily J Lelliott, Karen E Sheppard, David Goode, Rodney J Hicks, Tony Tiganis, Kaylene J Simpson, Ola Larsson, Carleen Cullinane, Vihandha O Wickramasinghe, Richard B Pearson, Grant A McArthur

## Abstract

Despite the success of therapies targeting oncogenes in cancer, clinical outcomes are limited by a residual disease that results in relapse. This residual disease is characterized by drug-induced adaptation, that in melanoma includes altered metabolism. Here, we examined how targeted therapy reprograms metabolism in BRAF-mutant melanoma cells using a genome-wide RNAi screen and global gene expression profiling. This systematic approach revealed post-transcriptional regulation of metabolism following BRAF inhibition, involving selective mRNA transport and translation. As proof of concept we demonstrate the RNA binding kinase UHMK1 interacts with mRNAs that encode metabolic proteins and selectively controls their transport and translation during adaptation to BRAF targeted therapy. Inactivation of UHMK1 improves metabolic response to BRAF targeted therapy and delays resistance to BRAF and MEK combination therapy *in vivo*. Our data support a model wherein post-transcriptional gene expression pathways regulate metabolic adaptation underpinning targeted therapy response and suggest inactivation of these pathways may delay disease relapse.

## Introduction

Clinical outcomes for cancer patients treated with oncogene targeted therapy are limited by residual disease that ultimately results in relapse. This residual disease is characterized by drug-induced cellular adaptation that precedes development of resistance. Maximum inhibition of oncogenic signalling has been the prevailing paradigm for improving antitumor responses to targeted therapies. For example, maximal suppression of BRAF-MEK signalling using combination therapy is current standard of care for BRAF mutant melanoma patients. Although this approach extended median survival to over 24 months from a historical base of less than 12 months (Larkin et al., 2014; Robert et al., 2015), the majority of patients still develop resistance and succumb to the disease. Targeting genetic features of drug resistant, relapsed disease has emerged as another paradigm to achieve more durable responses, however over 20 mechanisms of resistance have been identified in melanoma patients progressing on targeted therapy (Lim et al., 2017), revealing limitations in this approach. Prior to relapse, BRAF targeted therapy induces cellular adaptation that underlies residual disease (Menon et al., 2015; Rambow et al., 2018; Sharma et al., 2010; Su et al., 2017), and it has been proposed that non-mutational mechanisms underpinning this adaptability may provide new targets to improve clinical outcomes for patients.

Altered metabolism is a hallmark of cancer that has been intensely investigated over the last decade. How therapy reprograms metabolism and the role this plays during the adaptive response and development of resistance has received much less attention. In the setting of melanoma, we have previously shown that BRAF^V600^ inhibitor sensitivity correlates with glycolytic response in pre-clinical (Parmenter et al., 2014) and clinical studies (McArthur et al., 2012). BRAF inhibition (BRAFi) also renders BRAF^V600^ melanoma cells addicted to oxidative phosphorylation (OXPHOS) by releasing BRAF mediated inhibition of a MITF-PGC1A-OXPHOS pathway (Haq et al., 2013). This unleashes adaptive mitochondrial reprogramming, ultimately facilitating drug tolerance likely by compensating for suppressed glycolysis. Consistent with these observations, a “nutrient-starved” cell state emerges during the early drug adaptation phase following combined BRAF and MEK inhibition *in vivo*, and critically, cells appear to transition through this adaptive state as they acquire resistance (Rambow et al., 2018). Clinically, PGC1α (a biomarker for elevated OXPHOS) is induced in BRAF^V600^ melanoma patients treated with BRAFi, either alone (Haq et al., 2013) or in combination with MEK inhibitors (Gopal et al., 2014), whilst tumors that relapse following MAPK inhibitor treatment display an elevated mitochondrial biogenesis signature (Zhang et al., 2016). Together, these data suggest that maximal suppression of glycolysis and concurrent inhibition of adaptive mitochondrial metabolism may lead to improved outcomes to MAPK pathway targeted therapy by interfering with metabolic reprogramming underpinning drug-induced cellular adaptation. Notably, however, early results emerging from clinical trials of mitochondrial inhibitors such as biguanides have been largely disappointing (Kordes et al., 2015), and recent preclinical analyses support the concept that mechanisms underlying metabolic plasticity and adaptation may represent a more attractive therapeutic target (Hulea et al., 2018).

Here, we examined metabolic reprogramming in the therapeutic adaptation phase prior to acquired resistance using a genome-wide RNAi screen and global transcriptomic profiling. This systematic approach uncovered mRNA transport and translation pathways as regulators of metabolic response to BRAFi in BRAF^V600^ melanoma cells. Mechanistically, we demonstrate that metabolic response and adaptation is associated with selective mRNA transport and translation of metabolic proteins critical to BRAF inhibitor sensitivity and resistance, including glucose transporters and OXPHOS enzymes. This translational reprograming is mediated by the RNA binding kinase UHMK1 that is required for mitochondrial flexibility in response to BRAFi and controls the abundance of metabolic proteins through the export and translation of the mRNA that encode them. Importantly, genetic inactivation of UHMK1 increases sensitivity to BRAF and MEK combination therapy and delays resistance *in vivo*. Together, our data support a model wherein selective mRNA transport and translation contributes to metabolic adaptation underpinning therapy induced cancer cell plasticity, and suggests inhibition of this pathway may delay resistance to MAPK pathway targeted therapies.

## Results

### RNA binding, transport and translation pathways regulate metabolic response to inhibition of oncogenic BRAF signaling

To identify regulators of metabolic response following treatment with oncogene targeted therapy, we performed a genome wide RNAi glycolysis screen using BRAF^V600^ melanoma cells treated with the BRAF inhibitor (BRAFi) vemurafenib (Vem) as a paradigm (Figure 1A). Lactate is routinely used to measure glycolysis and can be readily detected in growth medium using a lactate dehydrogenase (LDH) enzyme based reaction. Cell number and viability were determined from nuclear DAPI staining using automated image analysis. For the screen, cells were first transfected with the human siGENOME SMARTPool library and subsequently treated with DMSO or a sub maximal dose of Vem (∼IC25)(Figure 1A). We chose a 48hr treatment which is within the window of metabolic adaptation following BRAFi, whereby maximal suppression of glycolysis (Parmenter et al., 2014) and adaptive mitochondrial reprogramming ((Haq et al., 2013; Zhang et al., 2016); Figure S1A) is observed. Notably, increased expression of SLC7A8 (LAT2), a biomarker of a drug tolerant “starved” melanoma state following BRAFi+MEKi *in vivo* (Rambow et al., 2018), was also observed (Figure S1B). Transfection of WM266.4 BRAF^V600^ cells with siRNA targeting polo-like kinase 1 (PLK1; death control) and pyruvate dehydrogenase kinase 1 (PDK1; glycolysis control) were used to define the dynamic range of the screening assays (Figure 1B), and notably, glycolysis was significantly more attenuated in Vem+siPDK1 cells compared to either Vem or siPDK1 alone, providing proof of principle for the major aim of the screen.

**Figure 1.**
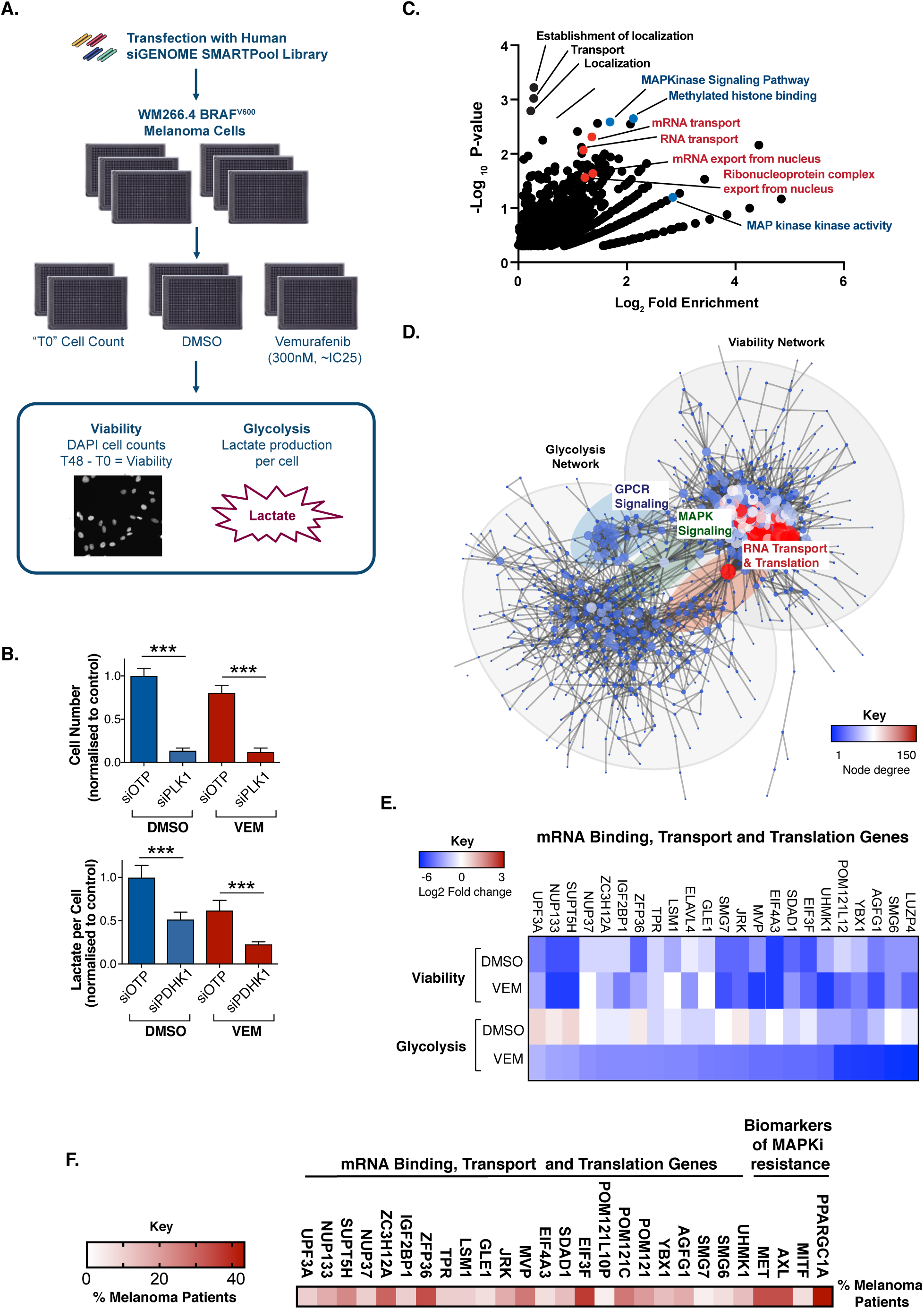
RNA binding, transport and translation pathways regulate metabolic response to BRAF inhibition. **A.** Schematic summarizing screen workflow (see methods). **B.** WM266.4 cells were transfected with the indicated siRNA and treated with DMSO or 300nM Vem for 48hr. Cell number was calculated using high content image analysis of DAPI stained cells (top panel) and growth media was collected for determination of lactate levels. Lactate absorbance values were normalized to cell number to determine lactate production per cell (bottom panel). Data is normalized to siOTP non-targeting (siOTP) transfected DMSO controls. Statistical significance was determined using a one-way ANOVA *** p > 0.001 (error bars = SEM, N=3). **C.** Functional annotation enrichment analysis was performed on 717 genes that enhanced the effects of Vem on lactate production (DMSO lactate per cell ratio < 0.5-fold change and Vem lactate per cell ratio > 0.5-fold change; see Table S1) using DAVID. Data is displayed as Log2 fold change versus - Log10 p-value. **D.** Network analysis was performed on 622 viability screen hits and 717 hits that enhanced the effects of Vem on lactate production using String (see Figure S1). Comparative network analysis was performed using Cytoscape, and hubs connecting the two networks are highlighted. **E.** Heat map displaying viability and lactate screening data for the indicated genes. **F.** Heatmap displaying percentage of melanoma patients with up-regulation of the indicated mRNA transport and translation genes on progression following treatment with MAPK pathway inhibitors (data sourced from (Hugo et al., 2015); see Table S4). See also Figure S1-2.

In the absence of drug, viability was impaired by depletion of 622 genes (Table S1) that formed a robust network (Figure S2A) enriched for regulators of cell cycle, translation and the ribosome (Figure S2B), processes previously shown to be critical for melanoma survival (Boussemart et al., 2014; Feng et al., 2015; Kardos et al., 2014). Glycolysis was reduced by depletion of 164 genes (Table S2), and as expected these genes were enriched with annotations associated with metabolism (Figure S2C and Table S2). To identify genes that regulate viability and glycolytic response to BRAFi, genes were grouped based on fold change data for each parameter in DMSO versus Vem treatment conditions (see supplementary information). This analysis identified 717 genes (Table S3) that were enriched for MAPK and GPCR signaling, and histone methylation, consistent with previous studies investigating BRAFi resistance (Figure 1C & Figure S2D)(Johannessen et al., 2013). However, surprisingly, the most striking feature of the gene set was RNA binding and transport, which was associated with 4 of the top 20 annotations ranked by P-value (Figure 1C & Table S3), with a total of 12 annotations associated with these pathways enriched in the dataset (Table S3). The identification of RNA binding and transport genes in our screen was particularly intriguing given these proteins are emerging as key determinants of gene expression programs activated in response to microenvironmental stress, including nutrient deprivation (El-Naggar and Sorensen, 2018). This group also included components of the EIF3 and EIF4F translation initiation complexes, and genes that regulate selective mRNA translation, thus also implicating mRNA translation in metabolic response to BRAFi. Notably, EIF4F has previously been reported as a nexus of resistance to MAPK pathway inhibitors in melanoma (Boussemart et al., 2014) thus further supporting performance of the screen. Comparative network analysis revealed 3 major hubs connect the viability and glycolysis networks; 1. GPCR signaling, 2. MAPK signaling, and 3. RNA transport and translation (Figure 1D), suggesting these pathways may coordinately regulate metabolic and viability responses to BRAFi. Consistently, 7 of the RNA transport and translation genes also enhanced the effects of Vem on viability (Figure 1E). The major findings of the screen were confirmed using a secondary de-convolution screen, whereby four individual siRNA duplexes were assessed to determine reproducibility of gene knockdown phenotypes. Notably, multiple RNA transport and translation genes were validated by 2 or more duplexes (33%; Figure S2E). We next assessed changes in expression of the RNA binding, transport and translation gene set using a published transcriptomic analysis of melanoma patients progressing after treatment with BRAF+/-MEK inhibitor treatment (Hugo et al., 2015). Strikingly, this analysis revealed that 18 out of 23 (78%) RNA transport and translation genes were upregulated in 10-36% of patients progressing on BRAF+/- MEK inhibitor treatment (Figure 1F & Table S4). By way of comparison, PGC1A was upregulated in 43% of patients in this dataset, whilst other previously documented biomarkers of acquired resistance to MAPK pathway inhibition in patients, c-MET and AXL, were upregulated in 33% of patients (Figure 1F). Viewed together, these large scale and unbiased analyses support a role for RNA binding, transport and translation pathways in regulation of metabolic response and viability following BRAFi.

### BRAFi induces transcriptional and translational reprogramming of metabolism in BRAF^V600^ melanoma cells

Given our functional screen suggested a role for post-transcriptional gene regulation pathways in metabolic reprogramming following BRAFi, we next assessed changes in mRNA abundance and translation efficiency by isolating total mRNA and mRNA bound to ribosomes using poly-ribosome (polysome) profiling (Figure 2A). Cell lysates were fractionated on a sucrose density gradient to isolate mRNA in sub-polysome (RNA-protein (mRNP) complexes and 40S, 60S, and 80S monomer peaks) or actively translating polysome (4 or more ribosomes) fractions (Gandin et al., 2014), and were analysed using RNA sequencing (RNA-seq). Of note, the number of ribosomes bound to mRNA is proportional to translation efficiency under most conditions. Global polysome profiles generated from DMSO treated A375 cells revealed a high basal rate of translation, and strikingly, this was potently suppressed by BRAFi at both 24 and 40hr (Figure 2B). Notably, this global inhibition of mRNA translation coincides with overt cellular adaptation (Figure S1) that presumably requires synthesis of new proteins, thus supporting the idea that selective mRNA processing and translation pathways play a role during the adaptive response to BRAFi.

**Figure 2.**
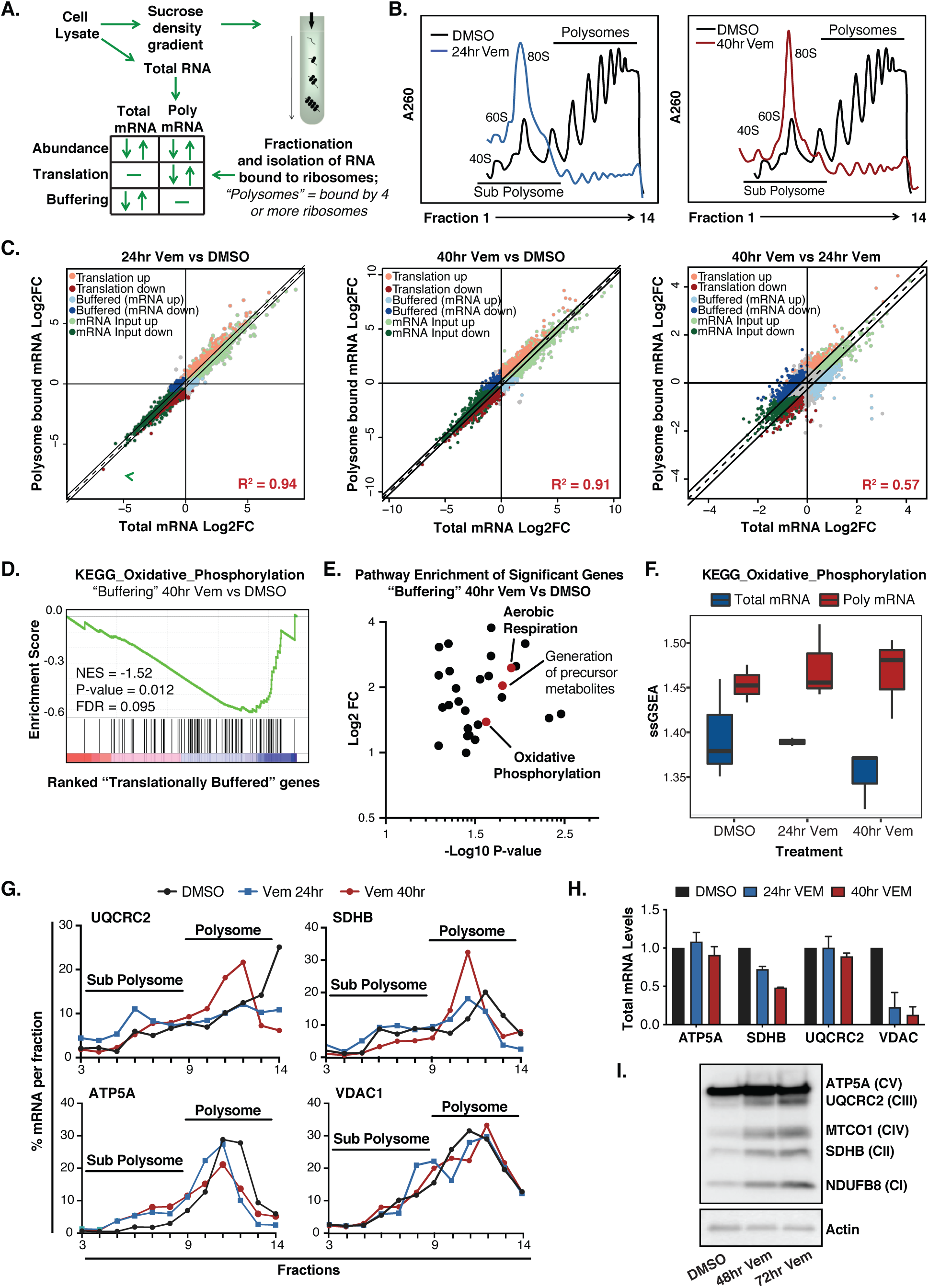
BRAFi induces transcriptional and translational reprogramming of metabolism in BRAF^V600^ melanoma cells. **A.** Schematic depicting the polysome profiling assay used to isolate total mRNA and polysome-bound mRNA (poly-mRNA) for RNA-seq analysis. Transcriptome-wide changes in different modes of gene expression were identified using anota2seq (mRNA abundance = changes in total and poly-mRNA; mRNA translation = changes in poly-mRNA only; translational buffering = changes in total mRNA and no change in poly-mRNA; see text for details). **B.** Polysome profiles of A375 cells treated with either DMSO or 1μM Vem for the indicated time on a 10-50% sucrose gradient (representative of N=3). **C.** Scatterplots of Log2 fold change (Log2FC) total mRNA vs polysome-bound (translated) mRNA in cells treated with DMSO or 1μM Vem for the indicated time. Different modes of gene expression identified by anota2seq are shown. **D.** Significantly enriched pathways for the different modes of gene expression were identified using GSEA (FDR < 0.1; see also Figure S3). GSEA plot demonstrating enrichment of the KEGG oxidative phosphorylation (OXPHOS) pathway is shown. **E.** Functional annotation enrichment analysis was performed on 579 significantly “buffered” genes (FDR < 0.1; Table S5) using DAVID (GO Biological Process and KEGG; P-value < 0.05; Table S7). **F.** Single sample GSEA (ssGSEA) pathway activity plot demonstrating translational buffering of the OXPHOS pathway. **G.** Distribution of mRNA encoding the indicated genes on a 10-50% sucrose gradient was determined using qRT-PCR following 1μM Vem treatment for the indicated time (representative of N=2). **H.** mRNA levels of the indicated genes was determined using qRT-PCR analysis of total mRNA samples (error bars = SEM, N=2). **I.** Whole cell lysates were analyzed by western blot for the indicated proteins following treatment with 1μM Vem for the indicated time (representative of N=3). See also Figure S3.

In order to identify transcriptome-wide changes in mRNA abundance and translation we used anota2seq (Figure 2A & Table S5)(Oertlin et al., 2019). Consistent with our previous studies (Parmenter et al., 2014), GSEA of changes in total mRNA levels revealed downregulation of multiple gene sets associated with the cell cycle and MYC transcription following 24hr of BRAFi, and these gene sets were further downregulated following 40hr treatment (Figure S3A and Table S6). In contrast, amongst the most significantly upregulated transcripts following 40hr BRAFi were biomarkers of the adaptive starved melanoma cell state identified *in vivo* (SLC7A8, CD36 and DLX5; Figure S3B)(Rambow et al., 2018). We next explored the global relationship between mRNA levels and mRNA translation efficiency during the drug treatment time course. Notably, although changes in total mRNA levels correlated strongly with changes in polysome association after 24hr and 40hr BRAFi compared to DMSO (R^2^=0.94, and 0.91 respectively), this relationship was less apparent when the 24hr and 40hr timepoints were compared (R^2^=0.57)(Figure 2C), indicating that changes in polysome associated mRNA cannot be solely explained by corresponding changes in mRNA abundance. These data indicate that mRNA transcription and processing is tightly coupled with mRNA translation efficiency within the early BRAFi response, however interestingly, this relationship appears to be uncoupled later during drug-induced adaptation (from 24-40hrs) indicating post-transcriptional modes of gene expression regulation. Analysis at the pathway level using GSEA also revealed differences between mRNA levels and mRNA translation 24hr and 40hrs post treatment, whereby the cell cycle and MYC pathways were the only significantly downregulated pathways in both datasets, and notably, decreases in translation efficiency of these pathways occurred later in the drug treatment at the 40hr time point (Figure S3A). Comparative analysis of total mRNA and polysome-associated mRNA levels identified genes with changes in total mRNA that were not reflected by a similar change in polysome-associated mRNA. These genes are termed “translationally buffered” (Figure 2A)(Oertlin et al., 2019), and indicate post-transcriptional mechanisms of gene regulation. GSEA of the translationally buffered gene set identified enrichment of multiple metabolic pathways, including pyrimidine metabolism and multiple OXPHOS gene sets (Figure 2D & S3C-D). Furthermore, functional annotation enrichment analysis of significant buffered genes (FDR < 0.1; Table S5) also revealed enrichment of OXPHOS and aerobic respiration (p < 0.05; Figure 2E & Table S7), further supporting post-transcriptional regulation of aerobic mitochondrial metabolism following BRAFi. Of note, these OXPHOS pathways were identified as “buffered (mRNA down)”, which corresponds to decreases in total mRNA levels and no change in polysome associated mRNA levels, as observed in single-sample GSEA pathway activity plots (Figure 2F & S3D). Importantly, discordance between translation efficiencies and total mRNA levels were validated for representative OXPHOS genes using qRT-PCR analysis of independently generated samples (Figure 2G-H), and consistent with the polysome profile analysis, OXPHOS protein levels corresponding to complexes I-IV were maintained or increased following BRAFi (Figure 2I). Indicating multiple modes of regulation for the OXPHOS pathway, no significant change in translation efficiency, total mRNA levels or protein levels, were observed for ATP5A (complex V). We also noted that MYC targets were enriched in the translational buffering dataset, potentially indicating that transcriptional downregulation of MYC targets may be uncoupled from mRNA translation. Because MYC-dependent regulation of glycolysis is a critical factor determining BRAFi sensitivity (Parmenter et al., 2014), we next explored adaptive translational buffering of MYC targets (Figure S3E) that relate to glycolysis, glucose transporter 1 (GLUT1) and hexokinase 2 (HK2). qRT-PCR analysis of GLUT1 and HK2 revealed total mRNA and polysome profiles consistent with translational buffering (Figure S3F-G), suggesting that any mRNA remaining in the cell following BRAFi will be translated at high efficiency. Interestingly, this data suggests that concordant inactivation of transcription and selective mRNA translation pathways may achieve more rapid and complete inactivation of glycolysis following BRAF targeted therapy, consistent with reduced lactate production in the orginal RNAi screen when expression of genes encoding regulators of mRNA processing were reduced.

Viewed collectively, these findings are consistent with our genome wide functional screen and support a role for selective post-transcriptional mRNA processing pathways in regulation of the proteome during early adaptive responses to BRAF. This includes key pathways implicated in metabolic reprogramming by BRAF and BRAFi sensitivity, MYC-driven glycolysis and oxidative mitochondrial metabolism.

### Depletion of the RNA binding kinase UHMK1 sensitizes BRAF^V600^ melanoma cells to BRAFi

Our systematic functional and transcriptomic approaches supported a role for selective RNA processing and translation pathways in metabolic response to BRAFi. Among the RNA processing proteins identified in our screen, U2AF homology motif (UHM) kinase 1 (UHMK1) was of most interest given it is the only known kinase to contain a classical RNA binding domain (the UHM domain), raising the hypothesis that it may function as a hub linking cell signaling and RNA processing. Moreover, UHMK1 regulates neuronal plasticity and adaptation via selective RNA transport and translation (Cambray et al., 2009; Pedraza et al., 2014) thus we hypothesized it may facilitate adaptive cellular reprogramming via RNA processing in the context of adaptation following BRAFi. We next investigated the role of UHMK1 in regulation of proliferative and metabolic responses to BRAFi in a panel of BRAF mutant melanoma cell lines (Figure 3 & S4). First, UHMK1 knockdown was confirmed using qRT-PCR and western blotting (Figure S4A). Because UHMK1 lacks a specific antibody for the endogenous human protein, we also confirmed increased levels of its key target p27, which is degraded following phosphorylation by UHMK1 (Boehm et al., 2002). siUHMK1+Vem treated cells showed more attenuated lactate production (Figure 3A), glucose utilization (Figure 3B), and extracellular acidification rates (ECAR; Figure 3C), when compared to BRAFi alone, indicating a reduction in glycolysis. A more marked reduction in cell number (Figure S4B) and cell proliferation (Figure 3D-E) was also observed in siUHMK1+Vem cells compared to Vem alone, and conversely, an increase in cell death was observed in 3 out of 4 cell lines (Figure 3F). Together these data confirm a role for UHMK1 in glycolytic, proliferative, and viability responses to BRAFi in BRAF^V600^ melanoma cells. Because UHMK1 kinase activity is required for regulation of mRNA processing in some contexts (Cambray et al., 2009; Manceau et al., 2008), we were next interested in exploring the role of UHMK1 kinase activity in BRAFi response. The kinase domain of UHMK1 shows limited homology to known kinases, however a K54A mutation in the putative active site extinguishes kinase activity (Maucuer et al., 1997). In order to assess UHMK1 kinase activity in BRAFi response, we first genetically inactivated UHMK1 using CRISPR-Cas9 and confirmed increased sensitivity of A375 cells to BRAFi (Figure 3G). Importantly this effect was rescued by expression of UHMK1-V5, but not the kinase dead K54A-V5 mutant (Figure 3G). UHMK1 expression and activity was confirmed using qRT-PCR and western blot analysis (Figure S4C). Viewed together, this data confirms a role for UHMK1 kinase activity in therapeutic responses to BRAF inhibition in melanoma cells.

**Figure 3.**
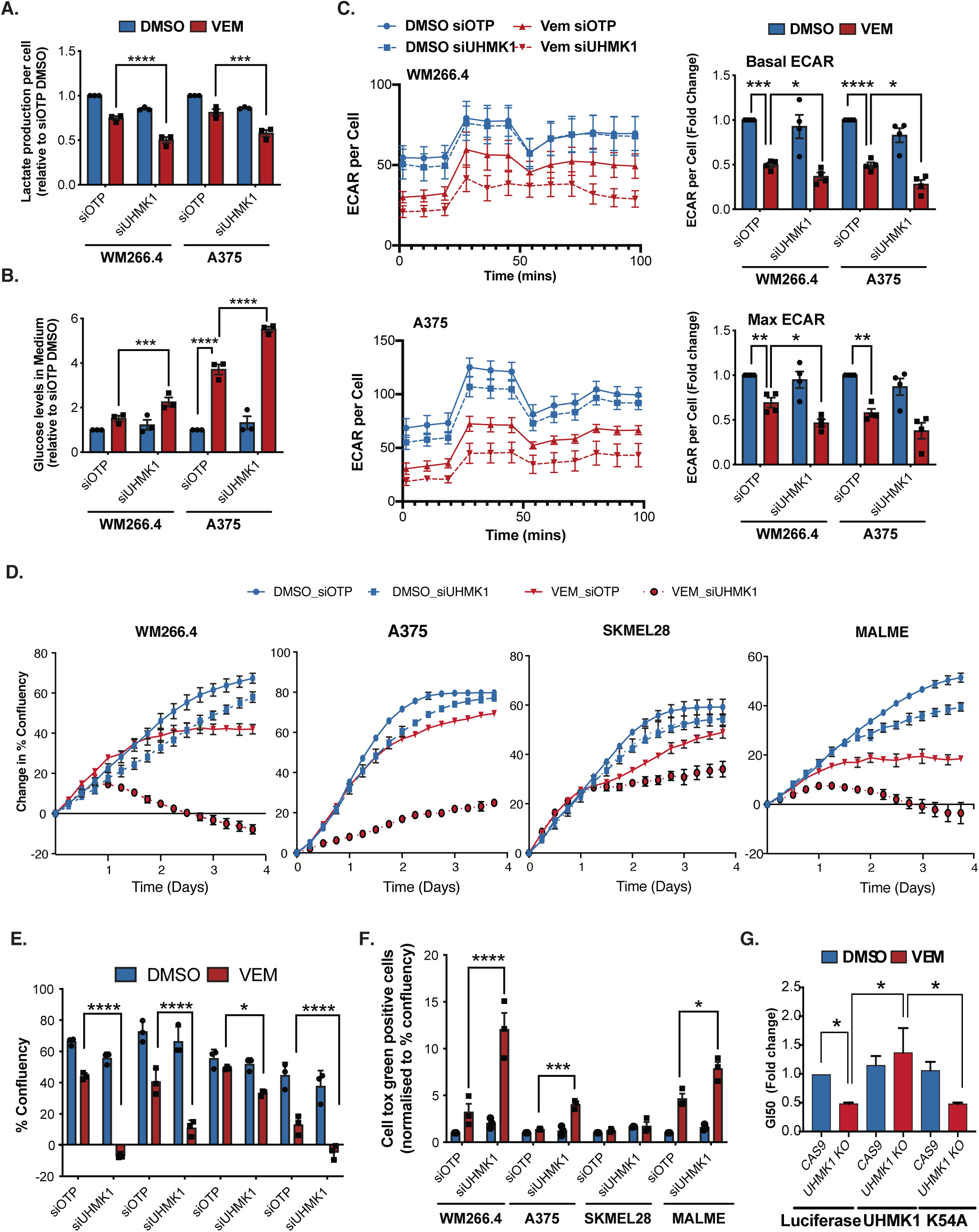
Depletion of RNA binding kinase UHMK1 sensitizes BRAF^V600^ melanoma cells to BRAFi. WM266.4 and A375 cells were transfected with the indicated siRNA and treated with DMSO or 300nM Vem for 48hr. Media was collected and lactate production **(A)** and glucose utilization **(B)** was determined. **C.** Extracellular acidification rate (ECAR) was determined using Seahorse Extracellular Flux Analysis and normalized to cell confluency (left panels). Basal ECAR was calculated from the third ECAR reading, and maximum (max) ECAR was calculated after treatment with the mitochondrial inhibitor oligomycin (fourth ECAR reading), and expressed as fold change relative to siOTP DMSO controls (error bars = SEM, N=3)(right panels). **D.** Cell proliferation was assessed in melanoma cells transfected with the indicated siRNA and treated with DMSO or 300nM Vem by monitoring confluency over time using an Incucyte automated microscope. Representative proliferation curves are shown. **E.** Average % confluency (normalized to T0) was calculated from proliferation data following 96hr treatment (error bars = SEM, N=3). **F.** Cell death was assessed in melanoma cells treated as in (E) using a Cell tox green cell death assay. Data is normalized to % confluency and expressed as fold change relative to siOTP DMSO controls (error bars = SEM, N=3). **G.** UHMK1 was genetically inactivated in A375 cells using CRISPR-Cas9, and a luciferase control, wild type UHMK1 or a K54A kinase dead mutant were ectopically expressed and sensitivity to Vem was assessed. Data is expressed as mean GI50 fold change relative to Cas9-Luciferase controls (error bars = SEM, N=4). Statistical significance was determined using a one-way ANOVA * p > 0.05, ** p > 0.01, *** p > 0.001, **** p > 0.0001. See also Figure S4.

### UHMK1 reprograms mitochondrial metabolism in response to BRAFi in BRAF^V600^ melanoma Cells

We next investigated whether UHMK1 can also promote adaptive reprogramming of mitochondrial metabolism in response to BRAFi in melanoma cells. Due to cell death after 72hr treatment with Vem+siUHMK1 (Figure 3F), we assessed cells after 48hr which immediately precedes overt mitochondrial reprogramming (Figure S1A). Analysis of oxygen consumption rates (OCR) using Seahorse extracellular flux analysis (Figure 4A) revealed only modest effects on basal and maximal OCR (Figure 4B & C) in Vem+siUHMK1 treated cells. However, significant reductions in spare respiratory capacity (Figure 4D) and ATP production (Figure 4E) were observed, indicating a reduced ability to respond to changes in energy demand and suggesting that UHMK1 can promote mitochondrial flexibility in response to BRAFi. Impaired mitochondrial metabolism in Vem+siUHMK1 treated cells was not associated with reduced mitochondrial number (Figure 4F), moreover only modest effects on PGC1A mRNA expression was observed (Figure 4G). We also assessed expression of mitochondrial transcription factor A (TFAM), another key regulator of mitochondrial biogenesis, and again saw no evidence of a role for UHMK1 in its expression. Instead, analysis of OXPHOS protein levels following Vem treatment revealed that increased expression of several OXPHOS proteins (NDUFB8, SDHB, UQCRC2, MTCO1) was UHMK1 dependent (Figure 4H). Together this data suggests that UHMK1 is required for regulation of oxidative metabolism following BRAFi by modulating OXPHOS protein expression independent of changes in the PGC1A- or TFAM-mitochondrial biogenesis transcription factor networks.

**Figure 4.**
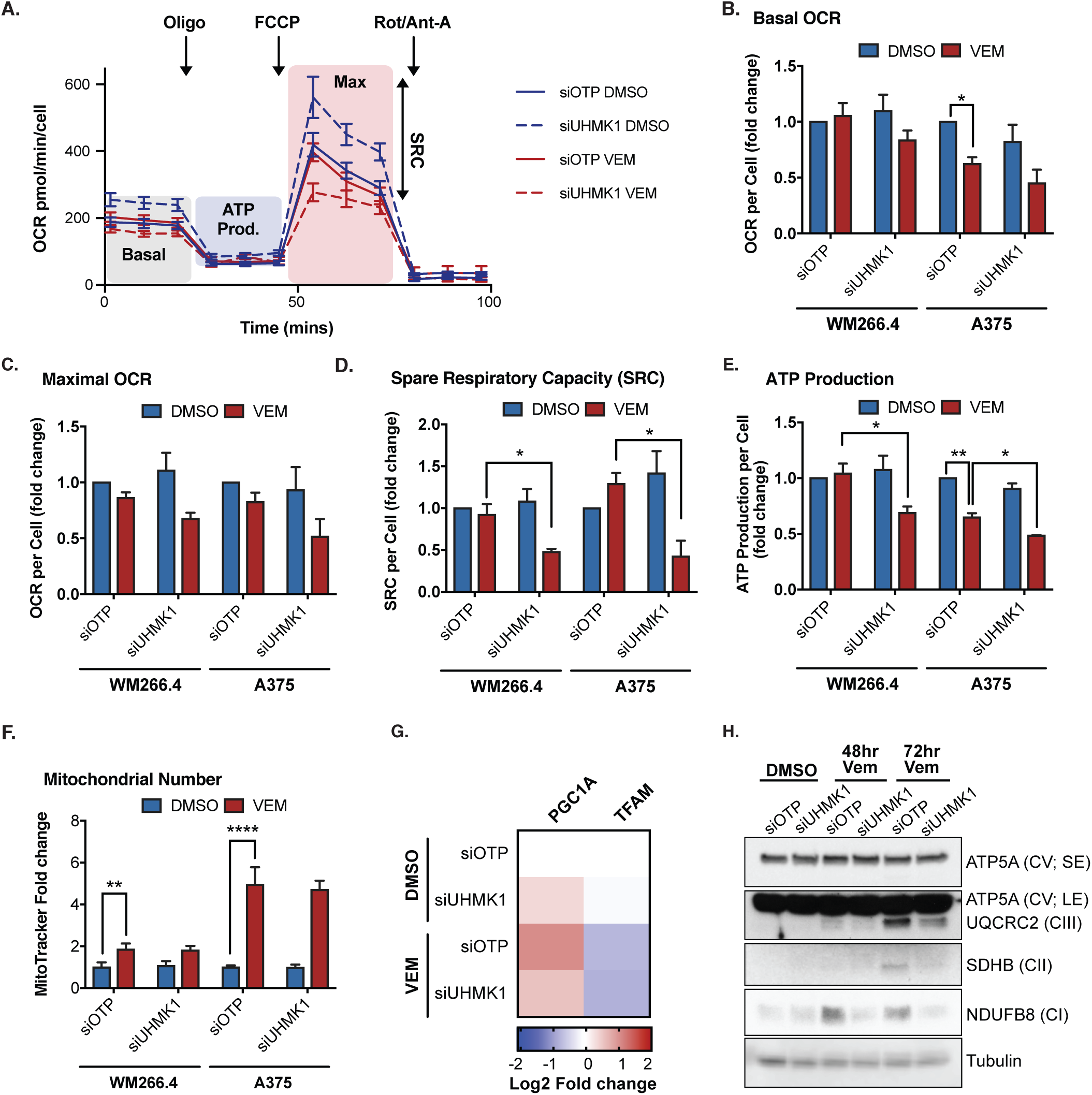
UHMK1 reprograms mitochondrial metabolism in response to BRAFi. WM266.4 and A375 cells were transfected with the indicated siRNA and treated with DMSO or 300nM Vem for 48hr. **A.** Oxygen consumption rate (OCR) was determined using Seahorse Extracellular Flux Analysis and representative profiles for WM266.4 cells are shown (Oligo = oligomycin; FCCP = Carbonyl cyanide-4-(trifluoromethoxy) phenylhydrazone; Rot/Ant-A = rotenone + antimycin-A; representative of N=4). Effect of gene knockdown and Vem treatment on basal OCR **(B)**, max OCR **(C)**, spare respiratory capacity (SRC) **(D)**, and ATP production **(E)** was determined following treatment with mitochondrial inhibitors as indicated in **(A)**. Data is normalized to cell number and expressed as fold change relative to siOTP DMSO controls (error bars = SEM, N=4). **F.** Mitochondrial number was determined using high content image analysis of Mitotracker stained melanoma cells treated as indicated (error bars = SEM, N=3). **G.** Effect of gene knockdown and Vem treatment on expression of the indicated genes was determined using q-RT-PCR. Data is expressed as Log2 fold change relative to siOTP DMSO controls. **H.** Whole cell lysates were analyzed by western blot analysis for the indicated proteins. Data is representative of N=3 (SE=short exposure; LE=long exposure). Statistical significance was determined using a one-way ANOVA * p > 0.05, ** p > 0.01, *** p > 0.001, **** p > 0.0001.

### UHMK1 binds to mRNA encoding metabolic proteins and regulates their nuclear-cytoplasmic transport in BRAF^V600^ melanoma cells adapting to BRAFi

In order to establish how UHMK1 regulates metabolic response to BRAFi, we next assessed its role in the mRNA expression pathway from transport to translation. The effect of Vem and UHMK1 knockdown on nuclear-cytoplasmic mRNA transport was first assessed using RNA fluorescence *in situ* hybridization (FISH) with an oligo(dT) probe which specifically binds to poly(A)^+^ pools of RNA (Figure 5A). In control conditions, the poly(A)^+^ signal was predominantly equal between the nucleus and cytoplasm (Figure 5A), however in contrast, depletion of the principal mRNA export factor NXF1 caused accumulation of the poly(A)^+^ signal in the nucleus (Figure 5A-B). Notably, nuclear accumulation of poly(A)^+^ mRNA was also observed in UHMK1 depleted cells, confirming a role for UHMK1 in mRNA transport in the context of melanoma cells. BRAFi also gave rise to a significant increase in the poly(A)^+^ nuclear to cytoplasm ratio (Figure 5B), however no further change was observed in the siUHMK1+Vem and siNXF1+Vem treated cells.

**Figure 5.**
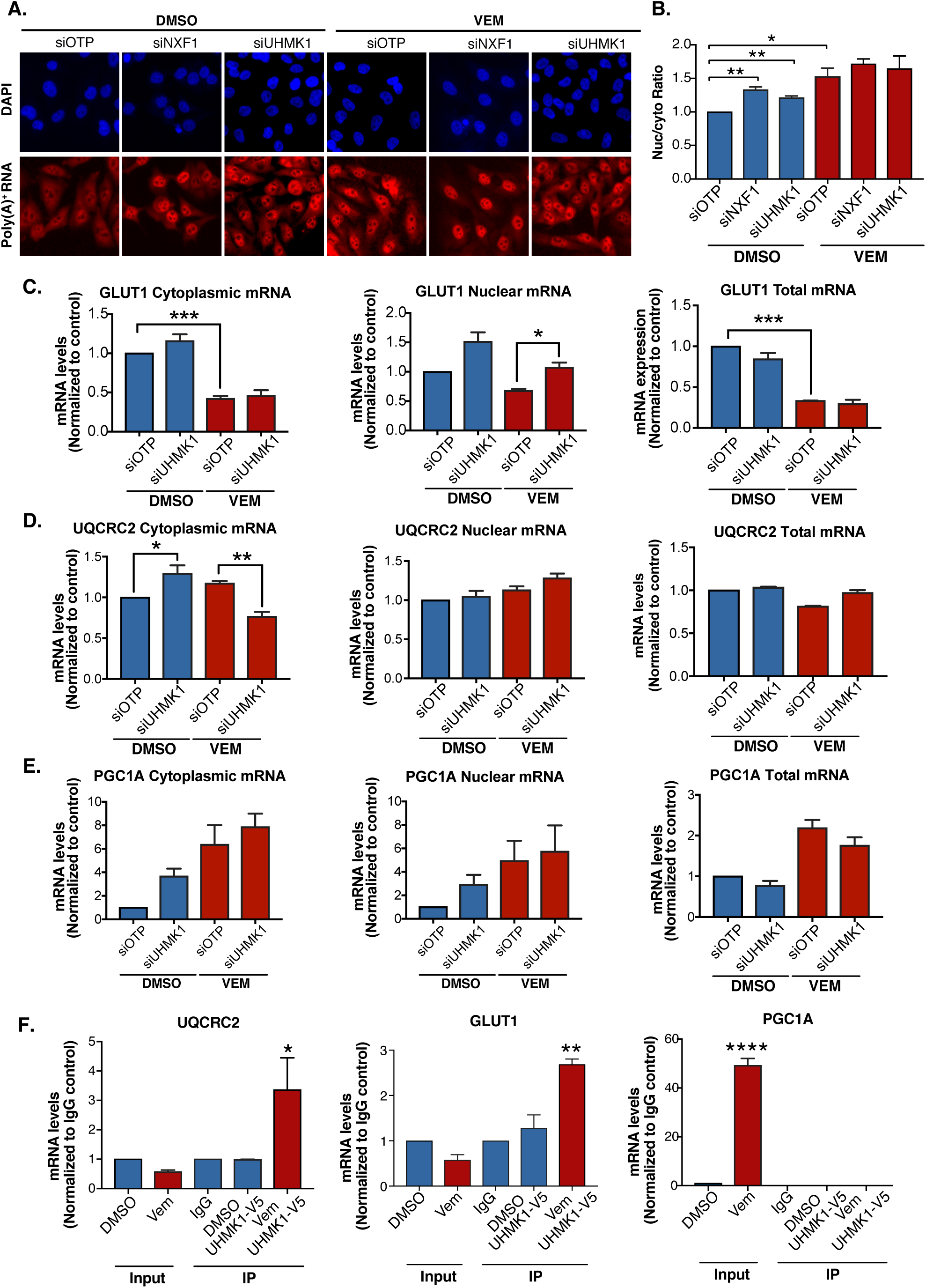
UHMK1 binds to mRNA encoding metabolic proteins and promotes selective mRNA transport in BRAF^V600^ melanoma cells adapting to BRAFi. A375 cells were transfected with the indicated siRNA and treated with DMSO or 1μM Vem for 48hr. **A.** RNA fluorescence *in situ* hybridization (FISH) using a poly(A)^+^ RNA specific probe in A375 cells treated as indicated (representative of N=3). **B.** The nuclear to cytoplasm ratio of poly(A)^+^ RNA was calculated using high content image analysis. Data is expressed as fold change relative to siOTP DMSO controls (error bars = SEM, N=3). **C-E.** Cell lysates were fractionated into nuclear and cytoplasmic pools of RNA and analyzed for the indicated genes using qRT-PCR. Whole cell lysates were used to assess total mRNA levels. **F.** RNA immunoprecipitation (RNA-IP) assays were performed in UHMK1-V5 expressing melanoma cells following treatment with DMSO or 1μM Vem for 48hr. The indicated mRNA transcripts were then analyzed using qRT-PCR. Statistical significance was determined using a one-way ANOVA * p > 0.05, ** p > 0.01, *** p > 0.001, **** p > 0.0001. See also Figure S5.

The more modest phenotype of UHMK1 compared to NXF1 depletion indicated a selective role for UHMK1 in mRNA transport. To extend these observations, we next assessed nuclear-cytoplasmic transport of specific mRNA transcripts using qRT-PCR analysis of nuclear and cytoplasmic mRNA pools generated from subcellular fractionation. The fractionation was verified by monitoring levels of mRNA known to be enriched within the nucleus (metastasis associated lung adenocarcinoma transcript 1; MALAT1) and cytoplasm (ribosomal protein S14; RPS14)(Figure S5A). We focused on GLUT1, HK2, and UQCRC2 that showed evidence of post-transcriptional regulation from our polysome profiling analysis. Notably, reduced cytoplasmic mRNA (UQCRC2) and increased nuclear mRNA (GLUT1 & HK2; Figure 5C-D & S5B) was observed specifically in the Vem+siUHMK1 treated cells, indicating UHMK1 depletion results in defects in GLUT1, HK2, and UQCRC2 mRNA transport following BRAFi. In contrast, analysis of PGC1A and ATP5A transcripts revealed no significant change in mRNA distribution (Figure 5E & S5B), consistent with no evidence of a role for post-transcriptional mechanisms or UHMK1 in their regulation from previous analyses (Figure 2&4). Together, these observations suggest that UHMK1 can selectively regulate nuclear-cytoplasmic mRNA transport in the context of therapeutic adaptation in BRAF^V600^ melanoma cells following BRAFi.

UHMK1 directly regulates localization and translation of specific mRNA transcripts by binding to mRNA (Cambray et al., 2009; Pedraza et al., 2014). To determine whether UHMK1 directly regulates mRNA encoding metabolic proteins following BRAF inhibition, we performed RNA immunoprecipitation (RNA-IP) assays using UHMK1-V5 expressing melanoma cells following DMSO or Vem treatment (Figure S5C). Strikingly, GLUT1, UQCRC2 and HK2 mRNA were not found in association with UHMK1-V5 in treatment naïve cells, however a significant increase in their association was observed following Vem treatment (Figure 5F & S5D). Further indicating specificity of the analysis and the pathway, no PGC1A mRNA could be detected in association with UHMK1 in any condition (Figure 5F). These data provide evidence that UHMK1 can directly regulate GLUT1, HK2, and UQCRC2 mRNA transport by physically associating with their mRNA, and strikingly, this association is induced by BRAFi.

### UHMK1 associates with polysomes and regulates selective translation of mRNA encoding metabolic proteins following BRAFi

To test the hypothesis that regulation of mRNA transport by UHMK1 selectively promotes translation of metabolic proteins following BRAFi, we next characterized the translational response of UHMK1 target genes following BRAFi. To achieve this we analyzed *de novo* synthesis of GLUT1 and OXPHOS proteins by giving a pulse with the methionine analogue L-azidohomoalanine (AHA), which is incorporated into all newly synthesized proteins (Figure 6Ai). This is followed by biotin labeling, streptavidin pull-down, and western blot analysis. Consistent with our polysome profile analysis, we observed a striking decrease in total AHA-labelled protein indicating a global inhibition of protein synthesis following 72hr Vem treatment (Figure 6Aii). In contrast, analysis of OXPHOS proteins following Vem treatment revealed a significant increase in *de novo* synthesis of UQCRC2 (Figure 6B-C), and significantly, increased synthesis of this OXPHOS protein was UHMK1 dependent. Again, supporting specificity of this pathway, no significant change in synthesis of ATP5A protein was observed (Figure 6B), consistent with polysome profiling of ATP5A mRNA (Figure 2D). Strikingly, we also observed that although GLUT1 protein synthesis was decreased following Vem treatment, this reduction was significantly more pronounced following UHMK1 knockdown (Figure 6B-C). This data suggests that UHMK1 depletion may cooperate with BRAFi to elicit a double-hit on the glycolysis pathway, whereby both GLUT1 mRNA transcription and translation is concurrently switched off. Linking these observations to UHMK1’s role in cellular responses to BRAFi, depletion of UQCRC2 and GLUT1 phenocopy UHMK1 knockdown whereby enhanced sensitivity to BRAFi was observed in cell proliferation assays (Figure S6A-C and (Parmenter et al., 2014)). However in contrast, no effect on Vem sensitivity was observed in the context of Vem+siATP5A treated cells (Figure S6C). Together, this data supports a model whereby UHMK1 regulates glycolysis and mitochondrial metabolism following BRAFi via mRNA transport and translation.

**Figure 6.**
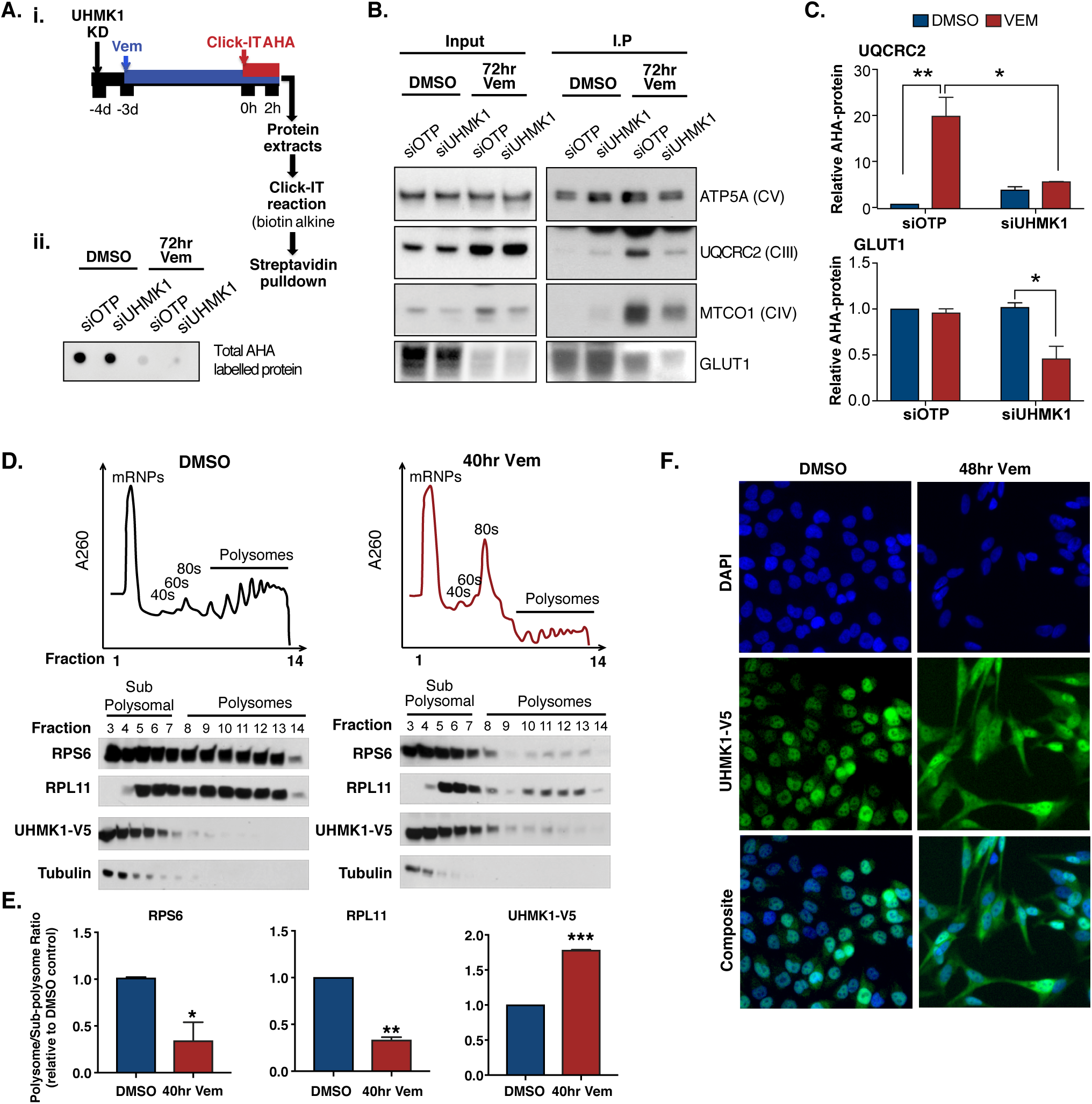
UHMK1 associates with polysomes and regulates selective translation of mRNA encoding metabolic proteins following BRAFi. **A.** Schematic depicting the AHA-based *de novo* protein synthesis assay **(i)** and dot blot **(ii)** showing total AHA labelled protein obtained from siOTP or siUHMK1 transfected cells following treatment with DMSO or 1μM Vem for 72hr. Data is representative of N=3. **B.** Protein lysates from input samples (left panel) and following streptavidin IP (right panel) were assessed using western blot analysis of the indicated proteins. **C.** Quantitation of AHA labelled protein shown in (B) (error bars = SEM, N=3). **D.** UHMK1-V5 expressing A375 cells were treated with DMSO or Vem for the indicated time, prior to polysome profiling. Representative profiles of 2 independent experiments are shown (top panel). Proteins were precipitated from the sucrose fractions and the indicated proteins were analysed using western blotting (bottom panel). **E.** Protein levels in sub polysome (fractions 3-8) vs polysome (fractions 9-14) fractions were calculated using densitometry, and sub polysome to polysome ratios were calculated (error bars = SEM, N=2). **F.** UHMK1 localization was assessed using high content image analysis of UHMK1-V5 expressing A375 cells treated with DMSO or 1μM Vem for the indicated time (representative of N=3). Statistical significance was determined using a one-way ANOVA * p > 0.05, ** p > 0.01, *** p > 0.001, **** p > 0.0001.

The mechanistic target of rapamycin (mTOR) pathway regulates translation and anabolic metabolism in response to nutrient supply (Ben-Sahra and Manning, 2017) and has been linked with a high OXPHOS phenotype in BRAF and MEK inhibitor resistance in melanoma cells (Gopal et al., 2014). Notably, although mTOR signaling was suppressed following BRAFi (Figure S7), no further changes to this pathway were observed following UHMK1 depletion, suggesting UHMK1 functions in an alternative pathway to modify mRNA translation during the early adaptation phase in cells treated with BRAFi. Differential association of mRNA processing and transport proteins with polysomes, and selective delivery of the transcripts they associate with, is another attractive hypothesis to explain translation of selective transcripts. To further explore the role of UHMK1 in adaptive programs following BRAF therapy, we precipitated proteins associated with polysomes using UHMK1-V5 expressing cells treated with DMSO or Vem (Figure 6D). As expected, small ribosomal protein RPS6 (a 40S ribosome component) was distributed in all fractions in control conditions, whilst large ribosomal protein RPL11 (an 80S ribosome component) was absent from early mRNP and 40S fractions. A significant reduction in the polysome to sub-polysome ratio was observed after Vem treatment (Figure 6E), consistent with global inhibition of translation (Figure 6D). Moreover, tubulin was restricted to sub-polysome fractions in both DMSO and Vem treated samples, further confirming specificity of the analysis (Figure 6D). In contrast, UHMK1-V5 protein was predominantly associated with sub polysome fractions in control conditions, however a redistribution of the protein to actively translating polysome fractions was observed following Vem treatment (Figure 6D-E). This data suggests that not only is UHMK1 recruited to polysomes in melanoma cells, but this association increases in response to BRAF therapy. Consistent with these observations, immunofluorescence analysis revealed a dramatic re-localization of UHMK1 from the nucleus to cytoplasm in cells treated with Vem (Figure 6F), and this was not associated with any change in UHMK1 protein levels (Figure S5Cii). Together this data supports a model whereby UHMK1 binds to mRNA and is translocated from the nucleus to the cytoplasm in response to BRAFi, where a proportion of the protein (∼13%) associates with polysomes and participates in selective regulation of mRNA translation.

### Genetic inactivation of UHMK1 sensitizes BRAF^V600^ melanoma cells to BRAF and MEK combination therapy *in vitro* and *in vivo*

We were next interested in testing the hypothesis that UHMK1 depletion would improve response and delay resistance following treatment with the current standard of care for BRAF^V600^ melanoma patients, a BRAF+MEK inhibitor combination. First, we performed cell proliferation assays and observed more attenuated proliferation in cells treated with the siUHMK1+BRAFi+MEKi triple combination compared to the BRAFi+MEKi combination alone (Figure 7A-B). To assess the role of UHMK1 in therapeutic response to BRAFi+MEKi *in vivo*, we implanted A375 cells expressing CAS9 or two independent UHMK1 gRNA into NOD scid interleukin 2 gamma chain null (NSG) mice (Figure 7C-D). Importantly, increased sensitivity to BRAFi+MEKi combination therapy was observed in mice implanted with both UHMK1 knock out cell lines compared with mice implanted with the control cell line (Figure 7E), culminating in a highly significant increase in overall survival (Figure 7F). Viewed together, this data confirms a role for the UHMK1 RNA processing pathway in MAPK pathway inhibitor responses in BRAF^V600^ melanoma cells both *in vitro* and *in vivo*.

**Figure 7.**
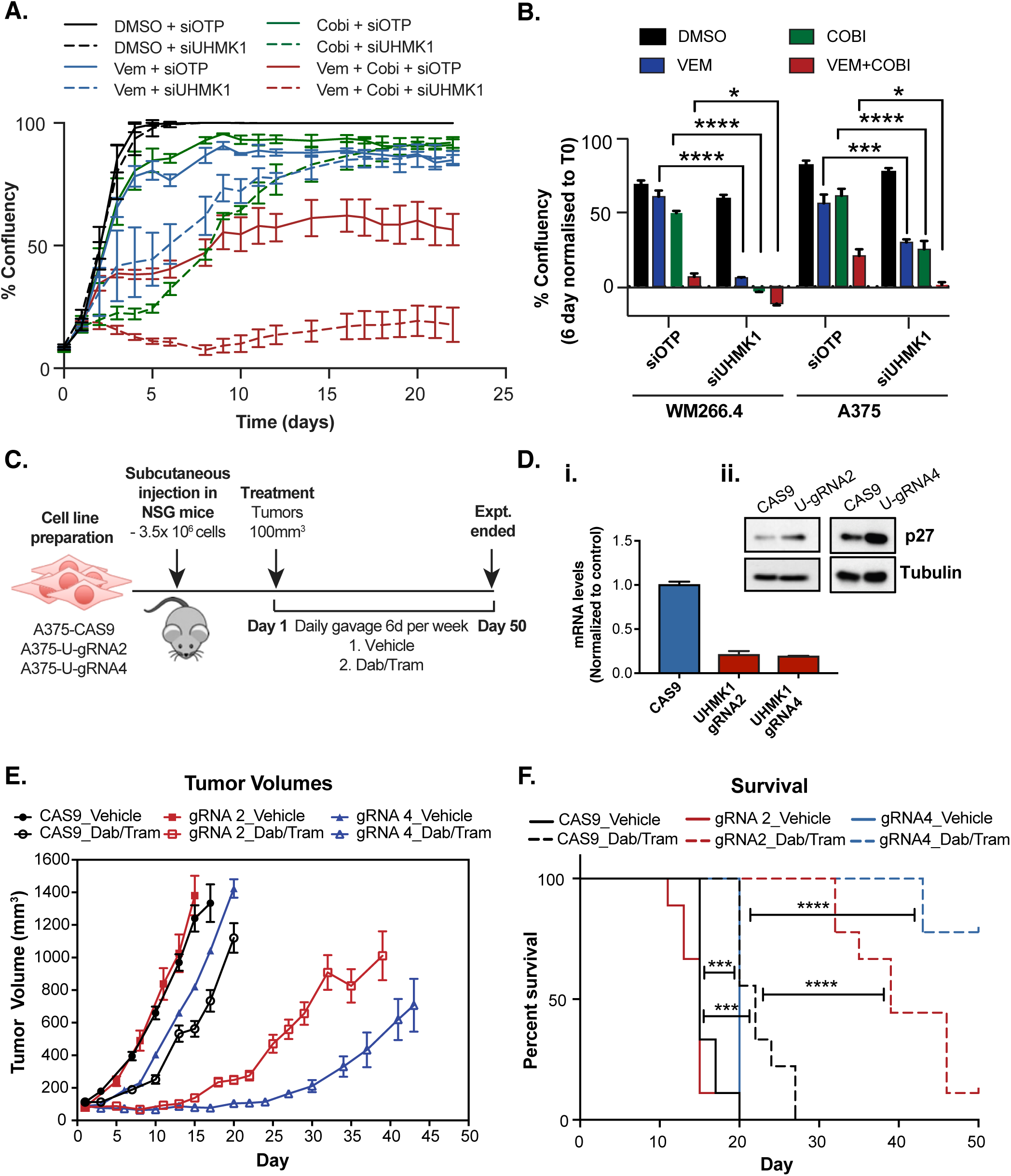
Genetic inactivation of UHMK1 sensitizes BRAF^V600^ melanoma cells to BRAF and MEK combination therapy *in vitro* and *in vivo*. Cell proliferation was assessed by monitoring confluency over time using an Incucyte automated microscope in melanoma cells transfected with the indicated siRNA and treated with DMSO, 300nM Vem, 10nM Cobi or Vem+Cobi. A representative proliferation curve **(A)** and average % confluency (normalized to T0) following 96hr treatment **(B)** is shown (error bars = SEM, N=3; top panel). **C.** Schematic of the *in vivo* drug sensitivity study. **D.** UHMK1 was genetically inactivated in A375 cells using CRISPR-Cas9 and UHMK1 KO was confirmed using qRT-PCR **(i)** and western blot analysis of UHMK1 target p27 **(ii). E.** Growth of A375-CAS9, A375-UHMK1-gRNA2 and A375-UHMK1-gRNA4 tumors treated with vehicle or dabrafenib and trametinib (Dab/Tram)(n=9 per group). **F.** Kaplan–Meier curve of data in **(E)** shows survival advantage where survival is defined as time to a tumor exceeding a volume of 1200mm^3^. ****P < 0.0001 by Log-rank (Mantel-Cox) test.

Altogether, our findings support a model wherein post-transcriptional gene expression pathways regulate metabolic adaptation underpinning targeted therapy response. As proof of concept, we demonstrate a role for UHMK1 in regulation of metabolic response and adaptation following BRAFi by controlling the abundance of metabolic proteins through the selective transport and translation of the mRNA that encode them. Importantly, inactivation of this pathway significantly improved survival following combined BRAF and MEK inhibition *in vivo*, suggesting inactivation of these pathways may delay disease relapse in melanoma patients.

## Discussion

Despite the success of therapies targeting oncogenes in cancer, clinical outcomes are limited by drug-induced adaptation and acquired resistance (Hugo et al., 2015). An emerging phenomenon observed following inhibition of oncogenic signaling in a range of cancers is suppression of glycolysis and adaptive mitochondrial reprogramming and enhanced reliance on oxidative metabolism (Baenke et al., 2015; Biancur et al., 2017; Caino et al., 2015; Ghosh et al., 2015; Haq et al., 2013; Hernandez-Davies et al., 2015; Kluza et al., 2012; Parmenter et al., 2014). In pancreatic cancer models, cells that survive genetic inactivation of KRAS^G12D^ display elevated mitochondrial metabolism, and treatment with an ATP synthase inhibitor delays relapse (Viale et al., 2014). Treatment of glioblastoma with PI3K inhibitors drives adaptive mitochondrial reprogramming that is associated with tumor cell invasion (Caino et al., 2015). Mitochondrial metabolism can also influence drug sensitivity, wherein analysis of melanoma patient samples linked mitochondrial gene expression signatures with intrinsic and adaptive resistance to BRAF and MEK inhibitors (Zhang et al., 2016). Inhibitors of oxidative metabolism, or the processes controlling adaptive mitochondrial reprogramming, are therefore attractive targets for combination therapy to circumvent acquired resistance before it can develop in a broad range of cancers. Here, we define a new mechanism of non-genetic drug adaptation whereby adaptive mitochondrial metabolism is regulated at the level of mRNA transport and translation and we identify the RNA binding kinase UHMK1 as central to this process. We propose inactivation of this pathway may represent a new strategy to interfere with adaptive metabolic reprogramming following oncogene targeted therapy, and delay resistance in melanoma patients.

mRNA translation has been implicated in responses to MAPK pathway inhibition and development of resistance in melanoma (Boussemart et al., 2014; Rapino et al., 2018). Here, we demonstrate that despite global suppression of translation during the early drug response phase, transcript selective translation reprograms mitochondrial metabolism through upregulation of OXPHOS proteins, and significantly, we show this is UHMK1 dependent. Importantly, upregulation of these mitochondrial proteins also occurs in melanoma patients progressing on BRAF and MEK targeted therapy (Zhang et al., 2016), linking these observations to resistance mechanisms in patients. Translational “buffering” (McManus et al., 2014) of glycolysis genes, whereby the rate of mRNA translation efficiency is maintained despite a decrease in total mRNA levels, also emerged from our analysis, and *de novo* protein synthesis assays revealed GLUT1 translation was maximally suppressed following UHMK1 depletion in combination with BRAFi. Viewed in the context of previous studies describing transcriptional repression of GLUT1 following BRAFi, this data supports a model whereby UHMK1 depletion cooperates with BRAFi to elicit a double-hit on the glycolysis pathway, whereby both GLUT1 mRNA transcription and translation is concurrently switched off. Consistent with these observations, Rapino *et al* (Rapino et al., 2018) have recently described codon-specific translational reprogramming of glycolytic metabolism in melanoma, in this case mediated by translational regulation of HIF1A by uridine 34 (U_34_) tRNA enzymes. Viewed together, these data suggest multiple mechanisms underpin transcript selective translational reprogramming of metabolism in cancer cells responding to oncogene targeted therapy. Interestingly, a recent report has also described translational reprogramming as a driver of phenotypic plasticity in the setting of melanoma cell invasion following glutamine deprivation (Falletta et al., 2017), indicating these pathways may also be part of a more general stress response activated by suppressed glycolytic metabolism when the BRAF oncogene is switched off.

However, in order for mRNA to be translated into protein it must first be exported from the nucleus and transported into the cytoplasm. This process is not always constitutive, as transcript selective RNA export pathways can regulate a range of adaptive biological processes including DNA repair, proliferation and cell survival (Wickramasinghe and Laskey, 2015). Interestingly, RNA binding proteins have recently been shown to regulate pro-oncogenic networks to control melanoma development (Cifdaloz et al., 2017), however their role in therapeutic response and oncogenic BRAF function has not been reported. Our work now implicates mRNA binding and transport as a driver of transcript selective translation following therapy, and we show that UHMK1 binds to mRNA encoding proteins relevant to metabolic response to MAPK pathway inhibitors and regulates their selective transport and translation. We suggest this mechanism allows cells to rapidly respond to cellular stimuli and stress such as nutrient deprivation. Interestingly, differential association of mRNA binding proteins with polysomes is one mechanism cells employ to rapidly regulate transcript selective translation (Aviner et al., 2017), and association of UHMK1 protein with polysome fractions following BRAFi is consistent with this concept. Moreover, a recent proteomic analysis of polysomes revealed 45% of all proteins identified were annotated as RNA binding, and a significant proportion of these were regulators of RNA transport and processing (Aviner et al., 2017). Notably, 78% of the RNA transport and translation gene set identified by the screen were upregulated in 10 - 36% of patients progressing on BRAF +/- MEK inhibitor treatment. Importantly, this is comparable to documented biomarkers of acquired resistance to MAPK pathway inhibition in melanoma patients, including PGC1A (43%), AXL (33%) and c-MET (33%)(Hugo et al., 2015), indicating relevance of these findings to human disease. Further analyses are now required to identify gene expression signatures associated with dysregulation of UHMK1 and other selective RNA binding and transport pathways in order to comprehensively assess their role in early adaptive reprogramming of metabolism and how this influences response to oncogene targeted therapies in cancer patients.

Our data suggests that UHMK1 inactivation does not influence selective RNA transport and translation through mTOR signaling, however UHMK1 activity is regulated by both AKT and ERK signaling in the context of growth factor stimulation (Lee and Kay, 2011). Further work is required to determine if UHMK1 may itself be a downstream target of the mTOR signaling network and function coordinately to control transcript selective translation in conditions of nutrient deprivation experienced by cancer cells following treatment with oncogene targeted therapy. Indeed, the eIF4E translation initiation factor, a component of the eIF4F translation complex subject to regulation by mTOR, has been implicated in mRNA processing, including nuclear-cytoplasmic mRNA export and transport (Bollmann et al., 2013; Culjkovic et al., 2005; Culjkovic-Kraljacic et al., 2012). Intriguingly, PGC1A expression and mitochondrial number remain unchanged by the BRAFi+siUHMK1 combination suggesting the UHMK1-RNA transport and translation pathway functions independently from the MITF-PGC1A-mitochondrial biogenesis pathway. Instead, a reduction in OXPHOS protein synthesis likely reduces the capacity of BRAFi+siUHMK1 treated cells to cope with glucose deprivation elicited by inhibition of BRAF signaling, a model supported by a reduction in spare metabolic capacity, and the ability of UQCRC2 knockdown to phenocopy UHMK1 depletion in combination with Vem.

Viewed collectively, our work supports a model wherein selective mRNA transport and translation is activated in response to therapeutic stress and contributes to metabolic reprogramming underpinning the adaptive therapeutic response. Our data demonstrate that the RNA binding kinase UHMK1 binds to mRNA encoding metabolic proteins critical to BRAFi response, and is required for their transport and translation following BRAFi. Inactivation of UHMK1 interferes with adaptive mitochondrial reprogramming following BRAFi, and critically, delays resistance and improves survival following combined BRAF and MEK inhibition *in vivo*. We propose that selective RNA transport and translation serves as a non-genetic mechanism of cancer cell adaptation and may provide a new target to interfere with drug adaptation and improve the efficacy of targeted therapies. We speculate this mechanism may also be relevant in broader oncogene driven cancer settings where responses to targeted therapies are blunted by phenotypic adaptation involving reprogrammed glycolysis and mitochondrial networks.

## Acknowledgements

We thank the following Peter MacCallum Cancer Centre core facilities: Victorian Centre for Functional Genomics (VCFG), Molecular Genomics, Flow Cytometry and the Centre for Advanced Microscopy and Histology. The VCFG (K.J.S.) is funded by the Australian Cancer Research Foundation (ACRF), the Australian Phenomics Network (APN) through funding from the Australian Government’s National Collaborative Research Infrastructure Strategy (NCRIS) program and the Peter MacCallum Cancer Centre Foundation. We thank Daniel Thomas, Jennii Luu, Kate Gould and Piyush Madhamshettiwar from the VCFG for technical and analytical assistance with the genome wide RNAi screen. We thank Dr Gisela Mir Arnau from Molecular Genomics for technical assistance with RNAseq. We also thank Rachael Walker and Susan Jackson from the Translational Research Lab and Alison Slater from the Molecular Oncology Lab for assisting in the *in vivo* studies. This work was supported by the Peter MacCallum Cancer Foundation and grants from National Health and Medical Research Council of Australia (#1053792 and #1106576) and the CASS Foundation (#8539). A.R, P.L and E.L were supported by doctoral scholarships from the University of Melbourne and Cancer Therapeutics Cooperative Research Centre.

## AUTHOR CONTRIBUTIONS

L.S, T.P, and G.M conceived and designed the project. L.S, T.P, K.J.S, and G.M designed experiments. L.S, T.P, M.K, E.K, J.K, T.W, and A.R conducted experiments. L.S, A.T, J.L and O.L performed data analysis. C.C and L.S designed and C.M, P.L, and E.L performed the *in vivo* experiments. V.W, R.P, T.T, K.E.S, C.C and R.J.H provided critical scientific input, protocols and/or reagents. L.S, V.W, R.P and G.M were involved in writing the manuscript, with all authors providing feedback.

