## Supplementary material for "Adaptive post-transcriptional reprogramming of metabolism limits response to targeted therapy in BRAF^V600^ melanoma": Figure S1

**Figure S1. Characterisation of metabolic adaptation following BRAFi in BRAFV600 melanoma cells.**

A.

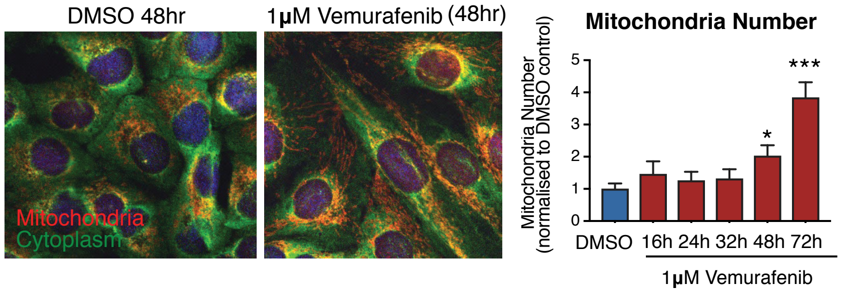


B.

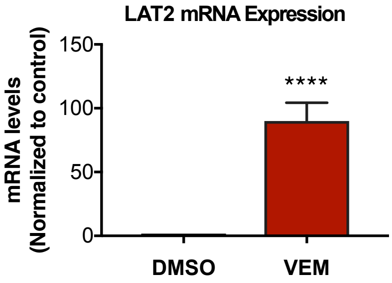
