## Supplementary material for "Adaptive post-transcriptional reprogramming of metabolism limits response to targeted therapy in BRAF^V600^ melanoma": Figure S3

**Figure S3. BRAFi induces transcriptional and translational reprogramming of metabolism in BRAF$^{V600}$ melanoma cells.**

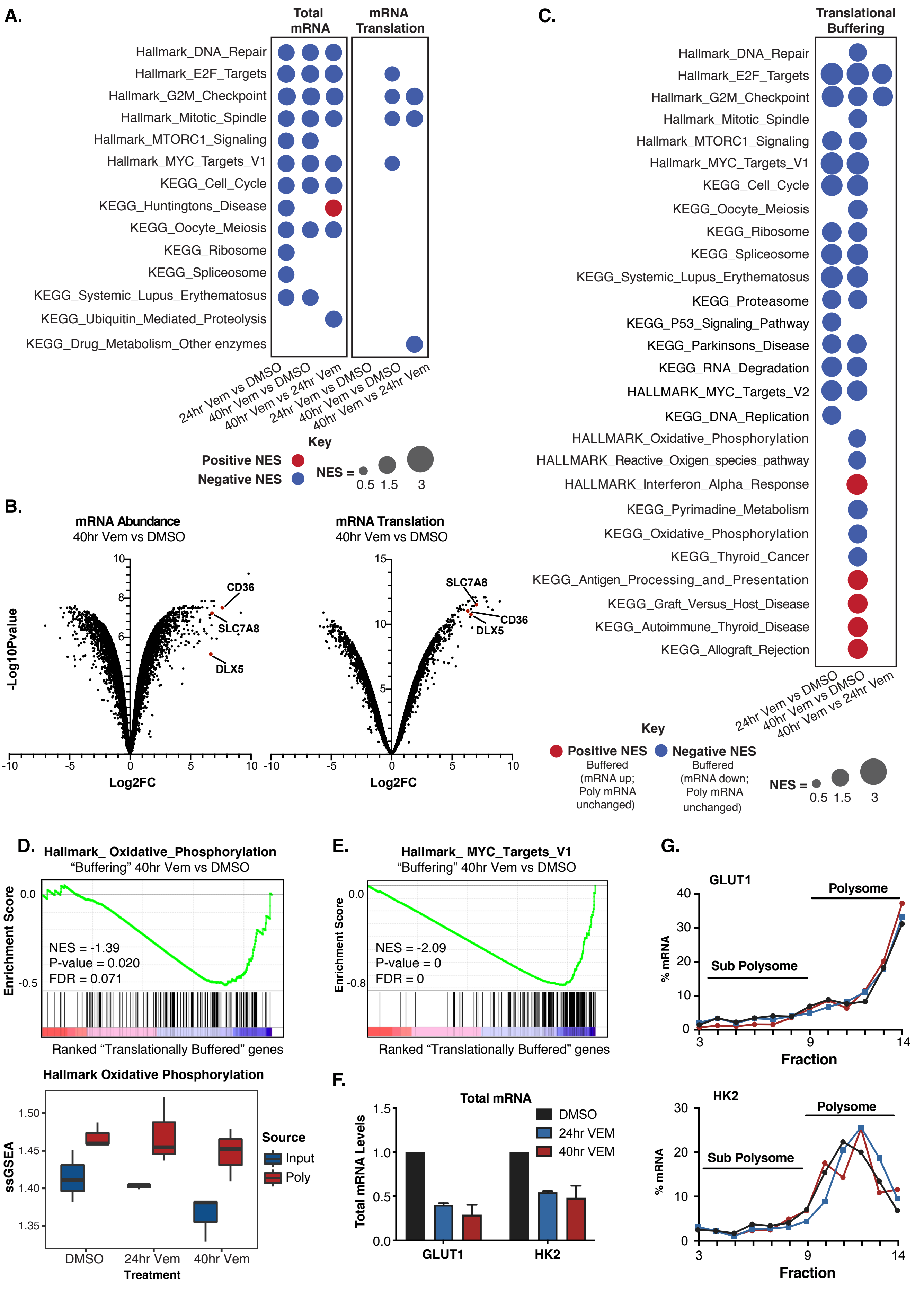
