## Supplementary figures and images for "Adaptive post-transcriptional reprogramming of metabolism limits response to targeted therapy in BRAF^V600^ melanoma"

### Figure S4

**Figure S4. UHMK1 depletion enhances the effects of BRAF inhibition in melanoma cells**

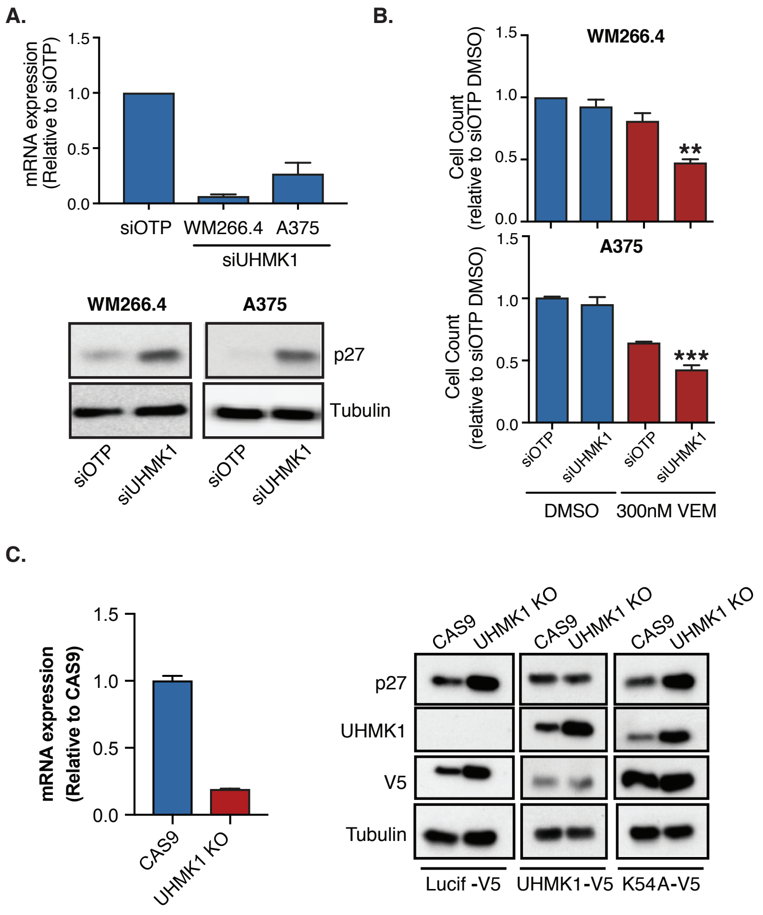

### Figure S6

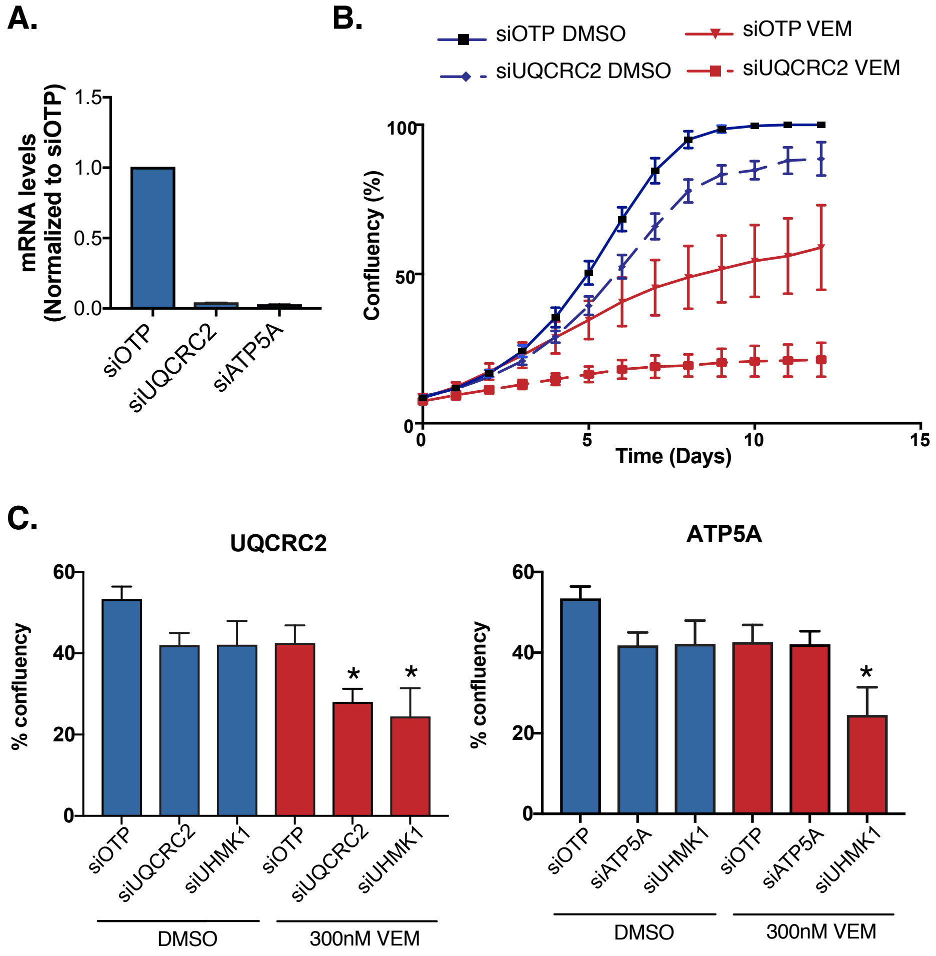
