## Supplementary material for "Adaptive post-transcriptional reprogramming of metabolism limits response to targeted therapy in BRAF^V600^ melanoma": Figure S5

**Figure S5. Analysis of the role of mRNA transport in UHMK1-dependent regulation of metabolism proteins following BRAF inhibition.**

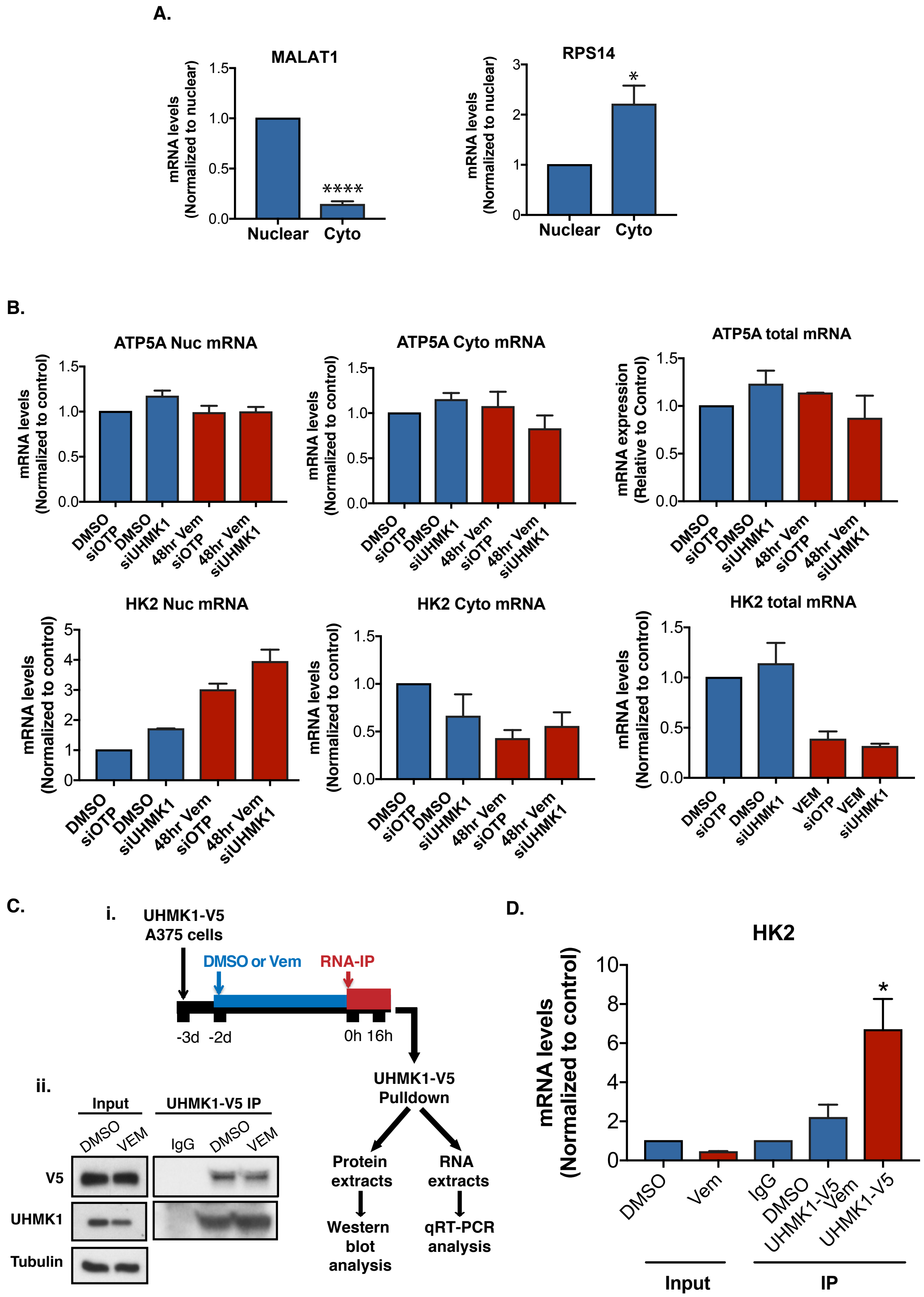
