## Supplementary material for "Adaptive post-transcriptional reprogramming of metabolism limits response to targeted therapy in BRAF^V600^ melanoma": Figure S7

**Figure S7. UHMK1 depletion modulates translation efficiency following BRAFi independent from the mTOR signaling pathway**

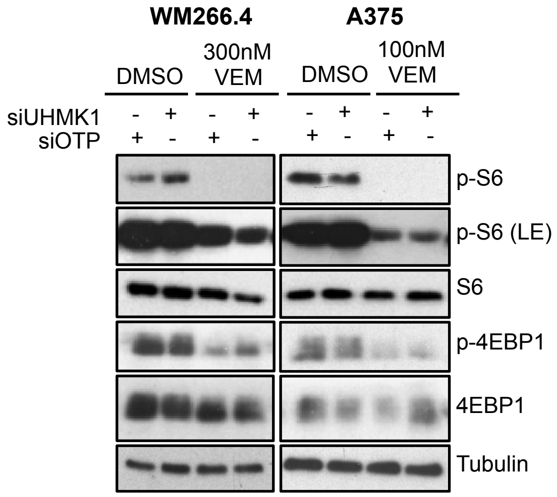
