## Supplementary Information for "Adaptive post-transcriptional reprogramming of metabolism limits response to targeted therapy in BRAF^V600^ melanoma"

**Supplementary Figure Legends**

**Figure S1. Functional characterization of adaptive metabolic reprogramming following BRAF inhibition in WM266.4 BRAF^V600^ melanoma cells**

**A.** A375 cells were treated with 1μM Vem for the indicated time and assessed for mitochondrial number using the mitochondrial stain MitoTracker. Representative confocal images are shown (left panel) and quantitation was performed using high content image analysis (right panel). **B.** Gene expression was determined using q-RT-PCR. Data is expressed as fold change relative to DMSO controls. Statistical significance was determined using a two tailed Students T test **** p > 0.0001.

**Figure S2. A genome wide screen for regulators of glycolytic responses to BRAF inhibition.**

**A.** Network analysis was performed on the 622 viability screen hits (DMSO ΔT48 > -1.5 Z-score; Table S1) using String. Enriched pathways were mapped onto the network as indicated. **B.** Functional annotation enrichment analysis was performed on the viability screen hits using DAVID. **C.** Functional annotation enrichment analysis was performed on the 164 glycolysis screen hits (DMSO lactate per cell ratio > 0.5-fold change; Table S2) using DAVID. **D.** Network analysis was performed on the 717 genes that enhanced the effects of Vem on lactate production (DMSO and Vem ΔT48 cell count > 0.3-fold change; DMSO lactate per cell ratio > 0.4-fold change; and Vem lactate per cell ratio < 0.5-fold change; Table S1) using String. Enriched pathways were mapped onto the network as indicated. **E.** Viability and glycolysis data from the secondary validation screen for genes associated with RNA binding, transport and translation. Individual siRNA duplexes (Du1-4) comprising each SMARTpool (SP) were arrayed into individual wells to confirm reproducibility of phenotypes. Data is expressed as Vem/DMSO fold change ratio.

**Figure S3. BRAFi induces transcriptional and translational reprogramming of metabolism in BRAF^V600^ melanoma cells**

**A.** Significantly enriched pathways for the indicated modes of gene expression were identified using GSEA (FDR < 0.1). Data is expressed as normalized enrichment score. **B.** Volcano plots of transcriptome-wide changes in total mRNA levels and polysome-bound mRNA (poly-mRNA; translation) identified using anota2seq (see methods for details; Table S5) following 40hr treatment with DMSO or 1μM Vem. **C.** Significantly enriched pathways for the translational buffering dataset (changes in total mRNA levels not reflected in changes in poly-mRNA levels) were identified using GSEA (FDR < 0.1). **D.** GSEA plot (top panel) and single sample GSEA (ssGSEA) pathway activity plot (bottom panel) demonstrating translational buffering of the Hallmark OXPHOS pathway. **E.** GSEA plot demonstrating enrichment of the Hallmark MYC targets (V1) gene set in the translational buffering data set. **F.** mRNA levels of the indicated genes was determined using qRT-PCR analysis of total mRNA samples (error bars = SEM, N=2). **G.** Distribution of mRNA encoding the indicated genes on a 10-50% sucrose gradient was determined using qRT-PCR following 1μM Vem treatment for the indicated time (representative of N=2).

**Figure S4.** **Depletion of the RNA binding kinase UHMK1 sensitizes BRAF^V600^ melanoma cells to BRAF inhibition.**

**A.** WM266.4 and A375 cells were transfected with the indicated siRNA and assessed for knock down efficiency using qRT-PCR (top panel) or western blot analysis (bottom panel). p27 protein levels were used as a biomarker of UHMK1 knockdown, as phosphorylation by UHMK1 targets p27 for degradation (see text for details). **B.** Effect of gene knockdown and Vem treatment on cell number calculated using high content image analysis of DAPI stained cells (error bars = SEM, N=3) **C.** UHMK1 was genetically inactivated using CRISPR-Cas9, and luciferase, UHMK1-V5 or a K54A-V5 kinase dead mutant were ectopically expressed. UHMK1 expression levels and activity (via p27-KIP levels) were assessed using qRT-PCR (left panel) and western blot analysis (right panel)(see text for details).

**Figure S5. Analysis of the role of RNA transport in UHMK1-dependent regulation of metabolism proteins following BRAF inhibition.**

A375 melanoma cells were transfected with the indicated siRNA and treated with DMSO or 1μM Vem as indicated for 48hrs. RNA was extracted from whole cell lysates or after nuclear and cytoplasmic fractionation. **A.** Nuclear and cytoplasm fractions were verified using qRT-PCR analysis of the indicated genes. Data is expressed as fold change relative to nuclear mRNA levels (error bars = SEM, N=3). **B.** Expression of the indicated genes was determined using qRT-PCR analysis (error bars = SEM, N=3). **C.** Schematic depicting the RNA immunoprecipitation (RNA-IP) assay (i) and western blot verifying UHMK1-V5 immunoprecipitation (ii). **D.** RNA was extracted from input and RNA-IP samples and analysed for the indicated genes using qRT-PCR (error bars = SEM, N=3). Statistical significance was determined using a two tailed T test, * p > 0.05, **** p > 0.0001.

**Figure S6. Depletion of UQCRC2, but not ATP5A, phenocopies UHMK1 knockdown following BRAF inhibition.**

**A.** A375 melanoma cells were transfected with the indicated siRNA and assessed for knock down efficiency using qRT-PCR. **B-C.** A375 melanoma cells were transfected with the indicated siRNA and treated with DMSO or 300nM Vem. Cell proliferation was assessed by monitoring confluency over time using an Incucyte automated microscope. A representative proliferation curve is shown **(B)** and average % confluency (normalised to T0) following 96hrs treatment was determined (error bars = SEM, N=3) **(C)**. Statistical significance was determined using a one-way ANOVA * p > 0.05.

**Figure S7. Analysis of the role of mTOR signaling and RNA transport in UHMK1-dependent regulation of mRNA encoding metabolic enzymes.**

WM266.4 and A375 melanoma cells were transfected with the indicated siRNA and treated with DMSO or Vem as indicated for 48hrs. Effect of gene knockdown and Vem treatment on protein levels was determined using western blot analysis.

**Methods**

**Cell lines and reagents**

Vem and its analog PLX4720 were provided by Plexxikon Inc. (Berkeley, CA, USA). Cobimetinib was purchased from Selleck Chemicals. All cell lines (WM266.4, A375, MALME, SKMEL28, HEK-293T) were purchased from the American Tissue Culture Collection (ATCC), and identity confirmed using STR profiling. All melanoma cell lines were maintained in RPMI 1640 containing 10% FBS, 2mM L-alanyl-L-glutamine in a 37°C humidified, 5% CO2 incubator. The BRAF and NRAS mutation status of all cell lines has been reported previously^1^. HEK-293T cells were cultured in DMEM containing 10% FBS, 2 mM L-alanyl-L-glutamine, in a 37°C humidified, 5% CO2 incubator.

**Genome wide RNAi glycolysis and viability screen**

The Dharmacon human siGENOME SMARTpool library (Dharmacon RNAi Technologies, Horizon Discovery) was used for the screen. The library was arrayed in 384-well format and screened within the Victorian Centre for Functional Genomics (VCFG, Peter MacCallum Cancer Centre, Australia). Each library plate was assayed in duplicate. All liquid handling steps were performed using a robotic BioTek 406 liquid handling platform, unless otherwise stated. All fixation and staining solutions were filtered (0.45μm filter) prior to use and plates were briefly centrifuged (500 x g for 30 sec) prior to all incubations.

*Screen method*

To perform the screen, a fresh vial of low passage WM266.4 cells (P8) were recovered and used for each individual batch of screening assay plates (58x library plates; 10-16 library plates screened each batch). For each library plate, 6x assay plates were required (2x no treatment cell number plates (T0), 2x 48hr control treated (0.1% DMSO), 2x 48hr drug treated (300nM Vem)). Cells were robotically seeded into columns 1-23 of black walled 384-well assay plates (450 cells/well;Corning) in 25μL growth media, and 25μL of media alone was added to column 24 for the lactate assay background control. Plates were pulse centrifuged (500 x g) and incubated for 10min on a level bench at room temperature (RT), then incubated overnight at 37°C in a Liconic STX200 automated microplate humidified incubator (37^o^C with 5% CO_2_). The transfection was performed 24hrs post cell seeding using a Caliper Sciclone ALH3000 liquid handling robot (Perkin Elmer, USA), RNAi MAX transfection lipid (Invitrogen, 0.03μL per well in 37.5μL) and siGENOME SMARTpool siRNA at a final concentration of 40nM. siOTP (D-001810-10-10) was used as the non-targeting control, siPLK1 (M-003290-01-0005) was used as a cell viability positive control, and siPDK1 (M-005019-00-0005) was used as a lactate assay positive control. Plates were pulse centrifuged (500 x g) and returned to the automated microplate incubator. 24hrs post transfection, transfection media was aspirated (z-height of 36) and replaced with 25μL of fresh media. Plates were pulse centrifuged to 500 x g and returned to incubator. 48hrs post transfection, media was aspirated (z-height of 36) from 4x assay plates and replaced with 25μL of fresh phenol-free RPMI media with 10% FBS and 2mM glutamine containing either vehicle (0.1% DMSO) or drug (300nM Vemurafenib). To generate “T0” cell number plates, 2x assay plates were fixed with 4% paraformaldehyde (PFA; Electron Microscopy Sciences, USA) and stained with DAPI DNA dye (1μg/mL) in PBS containing 10% triton X-100 (40μL per well) for 20mins. Plates were imaged on a Cellomics ArrayScan VTi automated microscope (Cellomics, Thermo Fisher Scientific, USA) using a 10x objective and 25 fields were captured per well. Image analysis and cell number calculation was performed using the Cellomics “Cell cycle” bioapplication. Optimal exposure time and object identification thresholds were identified for each individual batch of screening plates. To quantify lactate production per cell, media was collected from each assay plate 48hrs post drug treatment. Briefly, plates were centrifuged for 3mins (500 x g) and 10uL media was collected and transferred to a fresh plate using the Sciclone robot. Media was diluted 1:3 with PBS, mixed and stored at -80^o^C. Lactate concentration was determined using an L-lactate assay kit (Eton Biosciences) according to the manufacturers protocol. Briefly, 15μL of lactate reagent was added to 15μL media, mixed and incubated at 37^o^C in a CO_2_-free incubator for 45mins. The reaction was stopped through addition of acetic acid (0.5M) and absorbance (490nm) was read using a Cytation 3 Imaging Multi-Mode plate reader (Biotek). In parallel, cells were fixed and stained with DAPI and analyzed as described above. Background media absorbance was subtracted from experimental lactate absorbance values, converted to nM concentrations based on a lactate standard curve, and normalised to cell number to generate the parameter lactate production per cell.

*Screen analysis*

Data was expressed as fold change (FC) relative to the average of all siOTP non-targeting control wells included on each plate. Normalised sample values were averaged between replicated plates and hits were identified based on FC and robust z-score thresholds. Viability hits were identified based on cell number (T48 cell count; 0.3 FC, 723 hits) and change in cell number (ΔT48; T48 – T0 cell count; -1.5 Z-score, 622 hits) in vehicle treated plates (0.1% DMSO)(Table S1). Glycolysis hits were identified based on lactate production per cell (lactate absorbance/cell number) in vehicle treated plates (0.1% DMSO)(0.3 FC, 164 hits)(Table S2). Due to inaccuracies in lactate quantitation at low cell number, lactate data was filtered based on T48 cell count to remove genes with a FC < 0.3. To identify drug enhancers in the context of viability and lactate, genes were binned based on fold change data in control versus drug treated arms of the screen (DMSO and Vem ΔT48 cell count > 0.3; DMSO lactate per cell ratio > 0.4-fold change; and Vem lactate per cell ratio < 0.5-fold change; Table S3). Enrichment analysis for gene ontology (GO) terms (molecular function (MF) and biological process (BP)) and pathways (KEGG and Biocarta databases), was performed using DAVID (<https://david.ncifcrf.gov/>; Table S1-3). Protein interaction networks were identified using STRING (<https://string-db.org/>), and network data was visualized and analysed using Cytoscape.

*Secondary deconvolution validation screen*

To confirm the findings of the screen, we performed a secondary deconvolution validation screen, whereby each of the four individual siRNA duplexes were arrayed into individual wells to confirm reproducibility of phenotypes. The duplexes were screened at 25nM using the protocol described above. Duplexes were confirmed as “hits” if fold-change values for specific phenotypes were +/- 2-standard deviations of the median of non-targeting controls on each screening assay plate. DMSO/VEM ratios for viability and lactate were also calculated and used to define validated drug enhancement hits.

**Polysome profiling**

For polysome profiling, cells were pre-treated with 100µg/mL cycloheximide for 5 mins, washed with ice cold PBS containing 100µg/mL cycloheximide and lysed in a hypotonic lysis buffer (5 mM Tris-HCl (pH 7.5), 2.5mM MgCl2, 1.5 mM KCl, 100 µg/mL cycloheximide, 2mM DTT, 0.5% Triton X-100, and 0.5% sodium deoxycholate). Lysates were pre-cleared by centrifugation to remove nuclei, and the cytoplasm was collected and loaded onto a 10-40% linear sucrose density gradient (containing 20 mM Hepes-KOH (pH 7.6), 100 mM KCl, 5 mM MgCl2) and centrifuged at 36,000 rpm [SW40 Ti rotor (Beckman Coulter, inc)] for 2.15h at 4°C. Gradients were fractionated and collected (14 fractions per sample), and optical density was continuously recorded at 260nm using an ISCO Tris and UA-6 UV/VIS detector (Teledyne). Input RNA (20% lysate volume) was used to control for total amount of RNA per sample. RNA was isolated from sucrose fractions using phenol-chloroform extraction. For RNAseq, RNA pellets from fractions 9-14 (corresponding to polysome fractions) were pooled and further purified using RNeasy Mini Kits (QIAGEN), according to the manufacturer’s directions for RNA clean up. For analysis of individual mRNA transcripts using q-RT-PCR, RNA was isolated from individual fractions. For analysis of proteins, 10% trichloroacetic acid (final concentration) was used to precipitate protein from each fraction, and protein pellets were subsequently dissolved in SDS sample buffer and analysed using western blot analysis. The ratio of mRNA or protein associated with sub-polysome and polysome fractions was then calculated.

**RNA sequencing (RNAseq) and data analysis**

RNA quality and quantity was confirmed using Agilent Tapestation (Agilent Technologies), and all samples had an RNA integrity number (RIN) of 8.8 or higher. Approximately 1µg of RNA was used for library preparation using the TruSeq Stranded Total RNA Preparation Kit with Ribo-Zero Gold (Illumina). Briefly, ribosomal RNA (rRNA) was removed using biotinylated, target specific oligos and magnetic beads. The RNA was then fragmented using divalent cations under elevated temperature and reverse transcribed to cDNA with random primers. Indexed adaptors were then ligated and the library was amplified. Samples were then pooled and sequenced on a NextSeq500 (Illumina) high output flow cell to generate approximately 25 million single-end 75bp reads per sample. Library preparation and sequencing procedures were performed by the Molecular Genomics core facility at Peter MacCallum Cancer Centre.

Analysis of RNAseq data was performed using anota2seq as previously described^2^, with the following modifications. Genes with an average read count lower than 30 were removed from the analysis and data was normalized using the TMM-log2 approach and a batch effect (replicate number) was included in the models. Changes in polysome-associated mRNA (pool of efficiently translated mRNA i.e. mRNAs associated with 4 or more ribosomes) can be influenced by changes in corresponding total mRNA levels and/or be the result of changes in translational efficiency. Anota2seq allows to distinguish changes in amounts of polysome-associated mRNA that are independent of changes in corresponding total mRNA levels (regulation by mRNA translation) from fluctuations at the total mRNA level (regulation by e.g. transcription and/or mRNA stability). Furthermore, anota2seq can detect translational buffering which is another mode of regulation of gene expression where changes in polysome-associated mRNA and input total mRNA are also decoupled. In this case, polysome-associated mRNA levels (and protein levels) are preserved despite fluctuations in total mRNA levels. GSEA was performed on the gene lists generated by anota2seq using the preranked tool within the GSEA 3.0 software (Broad Institute). Genes were ranked based on Log2FC normalized for the adjusted p-value (Log2FCx1/adjp-val) and run against the Hallmark (V6.2) and KEGG (V6.2) gene sets. Gene sets with FDR < 0.1 were considered significant. Differentially expressed genes where filtered for fold change +/- 1.5 FC and adjusted p-values (Padj) < 0.1, and gene ontology enrichment analysis using the biological process and KEGG gene ontology sets was performed using DAVID. Gene ontologies with P-value < 0.05 were considered significant. Single-sample GSEA (ssGSEA)^4^ was performed using the GSVA R package^3^, which provides an enrichment score of the level of activity of gene sets in individual samples. The KEGG and Hallmark Oxidative Phosphorylation pathway gene sets used in the analyses were obtained from MSigDB c2 v6.2.

**siRNA-mediated gene knockdown**

Cells were forward-transfected with 40nM siGENOME SMARTpool siRNAs (Dharmacon) using 0.08 μl of Lipofectamine^TM^ RNAiMAX (Invitrogen) per 100ul of

transfection media per well, as per manufacturer’s directions. Briefly, RNAiMAX transfection lipid was diluted in OPTIMEM and equilibrated for 5mins, prior to complexing with siRNA for 20mins at RT. A non-targeting siOTP-NT siRNA was used as a control alongside siPLK1 as a technical control for cell viability . Media was changed 24h after transfection and plates were incubated at 37°C for indicated times and/or drug treated as described. Knockdown of selected genes were confirmed by qRT-PCR and Western immunoblotting.

**Plasmids and establishment of stable cell lines**

pLX304, pDONR-UHMK1, and pDONR-Luciferase were purchased from Addgene. pLX304-Luciferase and pLX304-UHMK1 expression vectors were generated using Gateway cloning (Invitrogen), following manufacturer’s directions. FuCas9Cherry was a gift from Dr. Marco Herold. The K54A kinase dead pLX340-UHMK1 mutant was generated using a QuickChange site-directed mutagenesis kit (Stratagene), as per manufacturer’s directions. HEK-293T cells were transfected with each plasmid along with the packaging plasmids pVSVG, pMDL and pRSV-rev by complexing with polyethylenimine (PEI). Virus generated by HEK-293T cells was supplemented with protamine sulfate (10 μg/ml), filtered, and transferred to melanoma cell lines four times for 12-16h. Virus-infected cells were selected with the appropriate antibiotic or by fluorescent activated cell sorting (FACS). All cell lines were verified using STR profiling and regularly tested for mycoplasma.

**CRISPR-CAS9 genome editing**

Synthetic guide RNAs (gRNA) targeting UHMK1 (gRNA-2 AACTGCTTGAGGGCGCCGGG and gRNA-4 CTTGCCGCCAGGAACCACCG) were designed using the Benchling online platform (<https://benchling.com/crispr>) and were synthesized by Sigma. A375-CAS9 stable cell lines were generated as described above and sorted for the top 30% expressing cells using FACS. CAS9 high expressing cells were transiently transfected with each gRNA (20nM diluted in 10mM TRIS-HCL pH7.5) and transactivator RNA (20nM diluted in 10mM TRIS-HCL pH7.5) using Dharmafect Duo transfection reagent. Cells were single cell sorted 72hrs post transfection into 96-well plates for single cell cloning. Clones were verified using sequence analysis of gDNA and q-RT-PCR analysis of mRNA.

**Metabolic assays**

For lactate production and glucose utilization assays, 15μL of phenol-free growth medium from treated cells was removed after pulse centrifugation (500xg). Growth media was diluted 1:3 with PBS and snap frozen at -80°C. Lactate levels were determined using an L-lactate assay kit (Eton Biosciences) as described above. Glucose levels were determined using a glucose fluorometric assay kit (BioVision) according to the manufacturers protocol. Absorbance (lactate) and fluorescence (glucose) were determined using a Cytation 3 Imaging Multi-Mode plate reader (Biotek). After the assay, cells were fixed and stained with DAPI DNA dye and cells were imaged using a Cellomics Arrayscan automated microscope. Image analysis and cell number calculation was performed using the Cellomics “Cell cycle” bioapplication (10x magnification; 16x fields), as described above. Lactate production and glucose utilization was normalised to cell number. To determine mitochondrial number, cells were labelled with MitoTracker (400nM for 30mins) according to the manufacturers protocol. Cells were fixed and stained with DAPI DNA dye and cells were imaged using a Cellomics Arrayscan automated microscope or on a Nikon C2 confocal microscope. Image analysis was performed using the Cellomics “Spot Detection” bioapplication (20x magnification; 16x fields).

**Extracellular Flux Analysis**

Extracellular flux analyses were performed on a Seahorse XF^e^24 or XF^e^96 Analyzer (Agilent, USA). For all assays, Flux Packs that contained the cell culture microplates, sensor cartridges and XF calibrant were used (Agilent 102416-100, 102340-100). Assay medium was prepared using Seahorse XF Base Medium DMEM (containing 5.5mM glucose, 2mM glutamine and 1mM sodium pyruvate, adjusted to pH 7.4 and kept at 37°C; Agilent 102353-100). Prior to cell seeding, Seahorse cell culture plates were coated with Corning Cell-Tak (438512) as per manufacturer’s directions. After the desired duration of gene knockdown and drug treatment, cell culture medium was removed and replaced with Seahorse XF medium and cells were equilibrated in a non-CO_2_ incubator for 1 hour prior to the assay. The XF Cell Mito Stress Test protocol was performed as per manufacturer’s directions, using oligomycin (1μM), FCCP (1μM) and rotenone/antimycin A (0.5μM). The assay was run with repeated cycles of 3min mix and 3min measurements following each drug injection with simultaneous measurement of OCR and ECAR. After the assay, cells were fixed and stained with DAPI DNA dye and imaged using a Cellomics Arrayscan automated microscope (10x magnification; 4x fields). Image analysis and cell number calculation was performed using the Cellomics “Cell cycle” bioapplication as described above. OCR and ECAR values were subsequently normalised to cell number.

**Dose response and proliferation assays**

Dose response assays were conducted in 96-well plates following 72hr drug treatments. Cells were fixed and permeabilized with methanol (MetOH), stained with DAPI nuclear dye and imaged using the Cellomics Arrayscan automated microscope (10x magnification; 16x fields). Image analysis and cell number calculation was performed using the Cellomics “Cell cycle” bioapplication as described above. Log[inhibitor] vs. response curves were generated by non-linear regression/curve fitting and GI50 concentrations (the concentration of drug required to reduce growth by 50%) were obtained as a measure of drug sensitivity. GI50s are displayed as mean ± SEM and statistical significance was determined using a Students *t*-test or one-way ANOVA (*p*<0.05). For proliferation assays, cells were plated at low density and treated with medium containing inhibitors 24hrs post seeding. Phase-contrast images were acquired and analysed daily using the IncuCyte (Essen Bioscience) continuous live-cell imaging and analysis system.

**RNA-FISH**

Cells were fixed and stained with a Cy3 labelled oligo(dT) primer (Sigma) and DAPI nuclear stain in black walled 96-well plates, as previously described ^5^. Plates were imaged on a Cellomics ArrayScan VTi automated microscope (Cellomics, Thermo Fisher Scientific, USA) using a 20x objective and 25 fields were captured per well. Image analysis and quantification was performed using the Cellomics “Nuclear translocation” bioapplication. A nuclear mask was generated from the DAPI channel and applied to the Cy3 oligo(dT) channel to calculate the average nuclear pixel intensity. A cytoplasmic mask 5 pixels wide was generated 1 pixel from the nuclear boundary in order to quantify the average cytoplasmic pixel intensity. The nuclear to cytoplasm ratio was calculated from these intensities.

**RNA fractionation**

Nuclear and cytoplasmic fractions were obtained by digitonin permeabilization of whole cells and centrifugation as previously reported ^5^. Briefly, cells were harvested in digitonin lysis buffer (50µg/mL digitonin (ICN Biomedicals), 100mM NaCl, 10mM Tris pH 8.0, protease inhibitors (Roche) and RNase inhibitor (Invitrogen)) and incubated on ice for 15mins. Following centrifugation at 1000xg for 5mins, the supernatant or cytoplasm fraction was separated from the pelleted nuclei, and RNA was isolated using RNeasy Mini Kits (QIAGEN), according to manufacturer’s directions.

**Protein-RNA immunoprecipitation**

Cells were harvested using a non-denaturing hypotonic buffer (5 mM Tris-HCl (pH 7.5), 2.5mM MgCl2, 1.5 mM KCl, 2mM DTT, 0.5% Triton X-100, 0.5% sodium deoxycholate, and RNAse inhibitor), and pre-cleared by centrifugation. After protein determination, samples were adjusted to 2mg and 10% volume was taken for protein and RNA input samples. Samples were incubated with antibody or IgG control at 4°C overnight under agitation. A/G Sepharose beads were blocked for 30 min in lysis buffer containing 10 mg/mL BSA and 0.1 mg/mL yeast total RNA, prior to incubation with lysates for 4 hr at 4°C under agitation. To elute protein, 50% of the sample was collected by boiling in 3x SDS sample buffer (187.5 mM Tris-HCl (pH 6.8), 6% w/v SDS, 30% glycerol, 150 mM DTT, 0.03% w/v bromophenol blue) and analysed using SDS-Page. To extract RNA, 1mL Trizol reagent was added to the remaining 50% of sample and RNA isolated following manufacturer’s protocol.

**RNA extraction and analysis of mRNA expression using Quantitative RealTime PCR (qRT-PCR)**

RNA was extracted using RNeasy Mini Kits (QIAGEN) and cDNA synthesis was performed using High Capacity cDNA Reverse Transcription Kits (Applied Biosystems), according to manufacturer’s directions. qRT-PCR was performed using Fast SYBR® Green PCR master mix using the primers listed below, on a Step One Plus Real time PCR system (Applied Biosystems). Data were processed using the comparative CT method, relative to the house keeping gene NONO. For RNA-IP experiments, data were analysed relative to the housekeeping gene β-actin. Changes in mRNA expression were expressed as fold change relative to assay controls, and analysed using a Students *t*-test or one-way ANOVA (*p*<0.05).

**qRT-PCR Primer Sequences:**

| **Gene** | **Direction** | **Sequence** |
| --- | --- | --- |
| ATP5A | Forward | CTTCGTTGCCACTTCCCAG |
|  | Reverse | CCTCCGGACTGGTTCTAGG |
| β-actin | Forward | CTTCCTGGGCATGGAGTC |
|  | Reverse | GGATGTCCACGTCACACTTC |
| c-Myc | Forward | GGACGACGAGACCTTCATCAA |
|  | Reverse | CCAGCTTCTCTGAGACGAGCTT |
| GLUT1 | Forward | TCTCTGTGGGCCTTTTCGTT |
|  | Reverse | CAGTTTCGAGAAGCCCATGAG |
| GLUT3 | Forward | GGTGGAAGTACGTTATTGTTGACTTATT |
|  | Reverse | GTTTGGCTAAAGGGTCTGAGATGT |
| HK2 | Forward | AAGGCAATAGGGCCTTAAAGTAGAG |
|  | Reverse | TTCGAGGCTGCAGTGAGCTA |
| HIF1α | Forward | TTTACCATGCCCCAGATTCAG |
|  | Reverse | GGTGAACTTTGTCTAGTGCTTCCA |
| MITF | Forward | CCGTCTCTCACTGGATTGGT |
|  | Reverse | TACTTGGTGGGGTTTTCGAG |
| NONO | Forward | CATCAAGGAGGCTCGTGAGAAG |
|  | Reverse | TGGTTGTGCAGCTCTTCCATCC |
| PGC1a | Forward | CTGCTAGCAAGTTTGCCTCA |
|  | Reverse | AGTGGTGCAGTGACCAATCA |
| SDHB | Forward | AGAAACTGGACGGGCTCTAC |
|  | Reverse | AACTGCAGGCCCCAGATATT |
| TFAM | Forward | TACCGAGGTGGTTTTCATCTG |
|  | Reverse | AACGCTGGGCAATTCTTCTA |
| UHMK1 | Forward | GCTGTTGATCTGTGGAGCCTA |
|  | Reverse | TCACCACTGCTTTACTGGCA |
| UQCRC2 | Forward | ATGTCCAAGCTGCCAAGAAC |
|  | Reverse | GGTGGCATGTAAGAACCAGC |

**Western immunoblotting**

Protein was extracted from cells using western solubilization buffer (WSB; 0.5mM EDTA, 20mM HEPES, 2% SDS), unless otherwise stated. Protein samples were subjected to SDS-PAGE analysis followed by western immunoblotting using the following antibodies: ERK (p44/42-MAPK) Cell Signaling Technology (CST) 9102, phospho-ERK (p44/42-MAPK; Thr202/Tyr204) CST 9101, 4EBP1 CST 9452, phospho-4EBP1 (Thr37/46) CST 2855, HIF1α AB2185, HK2 CST 1206, GLUT1 US Biologicals G3900-0J, PDHE1α Abnova H00005160-B01P, phospho-PDHE1α (Ser293) NB110-93479, MYC epitomics 1472-1, OXPHOS antibody cocktail AB110413, p27 BD transduction 610242, RPS6 CST 2217, phospho-RPS6 (Ser235/236) CST 2211, RPL11 Invitrogen 373000, S6-ribosomal protein CST 2217, phospho-S6 (Ser240/244) CST 2215, α-Tubulin Sigma T5168, UHMK1 Santa Cruz 393605, V5 CST 13202, YBX1 AB12148. For analysis of OXPHOS proteins using the OXPHOS antibody cocktail, lysates were boiled at 50°C for 10min, as per manufacturers directions.

***De novo* protein synthesis assay**

*De novo* protein synthesis assays were performed using the methionine analogue L-azidohomoalanine (AHA) as previously described^6^. Briefly, cells were starved for 30min in methionine-free RPMI media containing 10% dialyzed FBS and 0.2mM L-cysteine. Cells were given a pulse with AHA (100µM), and incubated for 2hrs at 37°C in a humidified 5% CO_2_ incubator. Cells were washed in PBS, harvested in lysis buffer (1% SDS in 50mM Tris-HCl pH 8.0) and boiled at 95°C or 50°C for OXPHOS proteins. Biotin labeling was performed using the Click-iT Protein Reaction Buffer kit (Invitrogen, C10276) as per manufacturer’s protocol, followed by streptavidin pull-down (Dynabeads M-280 streptavidin) and western blot analysis.

***In vivo* mouse experiments**

All animal studies were performed according to protocols approved by the Animal Ethics Committee of Peter MacCallum Cancer Centre and in accordance with the National Health and Medical Research Council Australian code for the care and use of animals for scientific purposes, 8^th^ Edition, 2013. 3.5x10^6^ A375-CAS9, A375-CAS9-UHMK1-gRNA2 or A375-CAS9-UHMK1-gRNA4 cells were prepared in 50% Matrigel and subcutaneously injected into the right flank of 6-7 week old female NOD-Scid interleukin 2 receptor gamma chain null (NSG) mice. Once tumours reached an average volume of 100mm^3^, mice were randomized into groups of 9 for therapy studies. Dabrafenib (30mg/kg in 0.5% HPMC and 0.2% Tween 80 in H_2_0) and trametinib (0.15mg/kg in 0.5% HPMC and 0.2% Tween 80 in H_2_0) were administered daily via oral gavage for 6 out of 7 days each week. Mice were euthanised when they met a tumor volume greater than or equal to 1200mm^3^.

**Statistical Analysis**

All statistical analyses were performed using GraphPad PRISM. Comparisons between two groups were analysed using the student’s t-test, and where more than two groups were compared, an analysis of variance (ANOVA) was performed, followed by the relevant multiple comparisons test. P < 0.05 was considered statistically significant. Un-clustered heatmaps were created in GraphPad PRISM.
