## Supplementary material for "Adaptive post-transcriptional reprogramming of metabolism limits response to targeted therapy in BRAF^V600^ melanoma": Figure S2

**Figure S2. A genome wide glycolysis screen for regulators of metabolic reprogramming in $BRAF^{V600}$ melanoma**

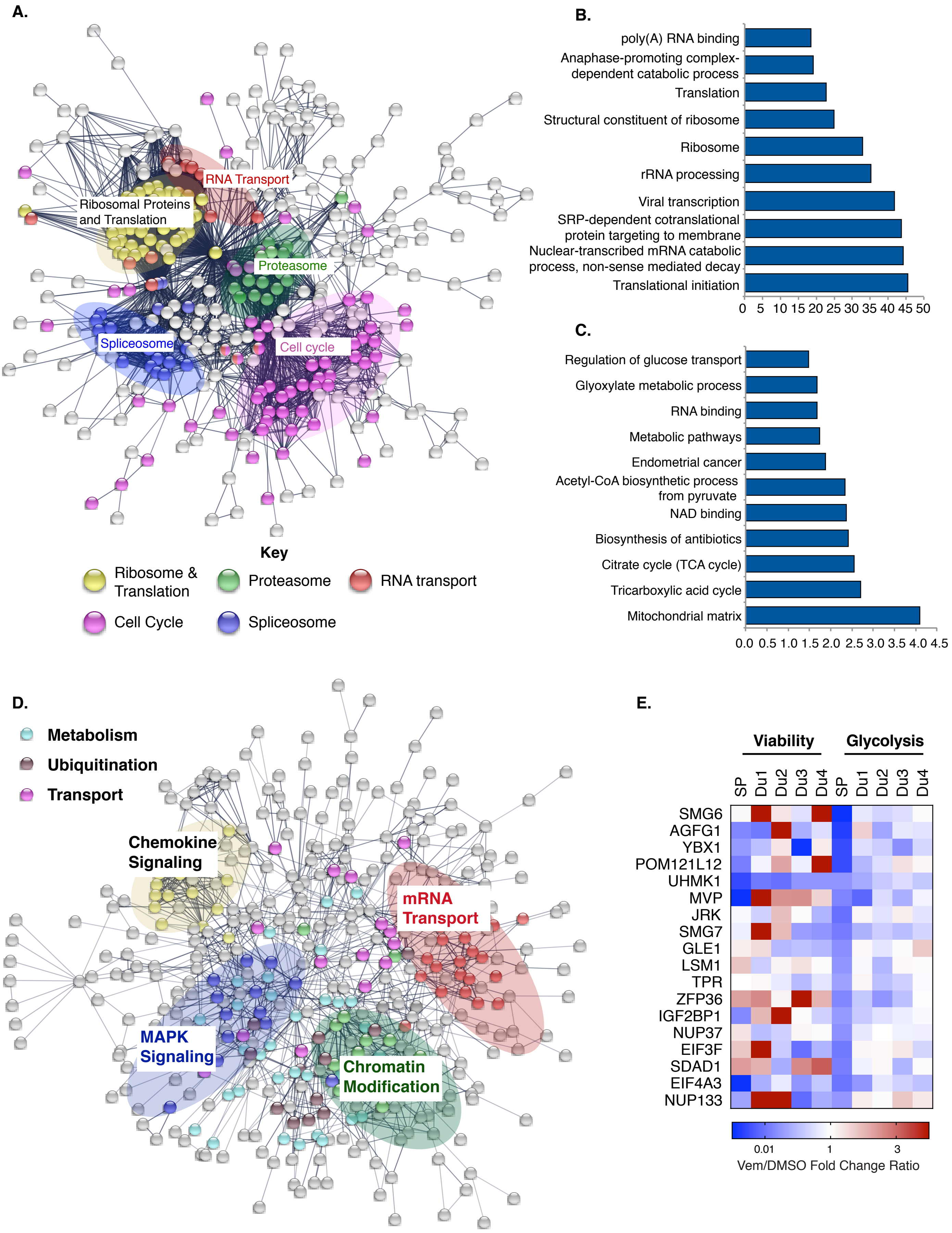
